# *Salmonella* lipopolysaccharide stimulates uptake of long-chain fatty acids in the small intestine

**DOI:** 10.64898/2026.05.19.726283

**Authors:** Eugene Koo, Gabriella Quinn, Cassie L. Behrendt, Brian Hassell, Katie N. Kang, Kelly A. Ruhn, Gonçalo Vale, Jeffrey G. McDonald, Wenhan Zhu, Sebastian E. Winter, Lora V. Hooper

## Abstract

Enteric bacterial pathogens profoundly alter intestinal physiology during infection, yet their effects on host lipid metabolism remain poorly understood. Using mass spectrometry lipidomics, we found that infection of the mouse intestine with the bacterial pathogen *Salmonella enterica* serovar Typhimurium (*S*. Typhimurium) stimulates uptake of long-chain fatty acids by small intestinal epithelial cells. This response coincided with increased expression of epithelial genes involved in lipid uptake and transport and required the long-chain fatty acid transporter CD36. Fatty acid uptake was triggered by *S*. Typhimurium lipopolysaccharide (LPS) and was impaired by bacterial mutations that alter LPS acyl chains. Mechanistically, *S*. Typhimurium induced lipid absorption through myeloid cell Toll-like receptor 4, a receptor for LPS. *Escherichia coli*, a related commensal bacterium, also induced intestinal lipid absorption through LPS, although to a lesser extent than *S*. Typhimurium. Finally, disruption of long-chain fatty acid absorption impaired host defense during bacterial stimulation, suggesting that bacteria-induced lipid uptake contributes to protection against enteric infection. Together, these findings identify LPS from Gram-negative intestinal bacteria as a key regulator of dietary lipid uptake by the intestinal epithelium.

**Significance Statement:** The intestinal lining absorbs nutrients while also detecting bacteria that enter with food, but how these two functions intersect during infection is poorly understood. *Salmonella* is a food-borne bacterial pathogen that infects humans. By studying mice, we found that *Salmonella* infection rapidly enhances absorption of dietary lipids by intestinal epithelial cells. This response was triggered when immune cells beneath the gut lining detected lipopolysaccharide, a molecule in the outer coat of many Gram-negative bacteria. In mice with defective lipid absorption, more *Salmonella* spread beyond the intestine, suggesting that lipid uptake contributes to host defense. These findings reveal a previously unappreciated link between microbial sensing, nutrient absorption, and defense against intestinal infection.

## Introduction

The gut microbiota plays a central role in shaping host physiology, with growing evidence indicating its profound influence on nutrient metabolism and energy balance. Among its best-characterized functions is the capacity to enhance digestion of complex carbohydrates, enabling the liberation and absorption of monosaccharides and short-chain fatty acids (1). In parallel, microbial signals strongly influence intestinal lipid absorption and systemic lipid metabolism (2–4). Previous studies have shown that the microbiota engages host-intrinsic pathways in epithelial cells to promote the transcription of lipid metabolism genes, thereby facilitating the uptake and processing of dietary lipids (5–7). These microbiota-driven effects on lipid handling have been linked to circadian regulation (5), histone modification (7), and long noncoding RNA activity (6), and require innate immune signaling via myeloid cells and group 3 innate lymphoid cells (5–7). Chronic exposure to microbial signals thus programs epithelial lipid uptake in a manner that contributes to long-term energy storage and the development of obesity.

By contrast, less is known about how enteric pathogens influence host lipid metabolism. While intestinal pathogens such as *Salmonella enterica* serovar Typhimurium (*S*. Typhimurium) have a well-recognized ability to trigger inflammation and disrupt barrier function (8), their impact on host nutrient handling has not been fully explored. Although enteric pathogens are known to alter carbohydrate availability and utilization during infection (9), it remains unclear whether and how they influence lipid absorption and utilization. Given that infection induces a robust innate immune response and alters epithelial cell gene expression and function (10–12), we reasoned that acute exposure to a pathogen might impact host lipid metabolism.

Here, we show that infection with *S*. Typhimurium rapidly enhances long-chain fatty acid absorption in the small intestine, coincident with the induction of epithelial transcriptional programs involved in lipid uptake and transport. This response is driven by bacterial lipopolysaccharide (LPS), depends on the acylation state of LPS, and requires Toll-like receptor 4 (TLR4) signaling in subepithelial myeloid cells. We further find that the related commensal bacterium *Escherichia coli* similarly promotes lipid absorption through an LPS-dependent mechanism. Together, these findings identify bacterial LPS as an acute regulator of host lipid metabolism and indicate that both pathogens and commensals can engage shared innate immune pathways to influence nutrient handling at the intestinal barrier.

## Results

### *S.* Typhimurium infection promotes lipid uptake and metabolic reprogramming in small intestinal epithelial cells

To investigate whether *S*. Typhimurium infection alters intestinal lipid metabolism, we reanalyzed a published single-cell RNA-seq dataset of intestinal epithelial cells (IECs) from infected and uninfected mice (12). This analysis revealed increased expression of genes involved in lipid metabolism and transport in IECs following infection (Fig. 1A–C). Gene set enrichment analysis of differentially expressed transcripts showed that lipid metabolism pathways were prominently represented among genes upregulated during infection (Fig. 1B), accounting for more than 10% of enriched pathways (37 of 313). Notably, genes involved in multiple stages of lipid trafficking, including lipid import, intracellular transport, and export, were increased in IECs from infected mice (Fig. 1C).

**Figure 1.**
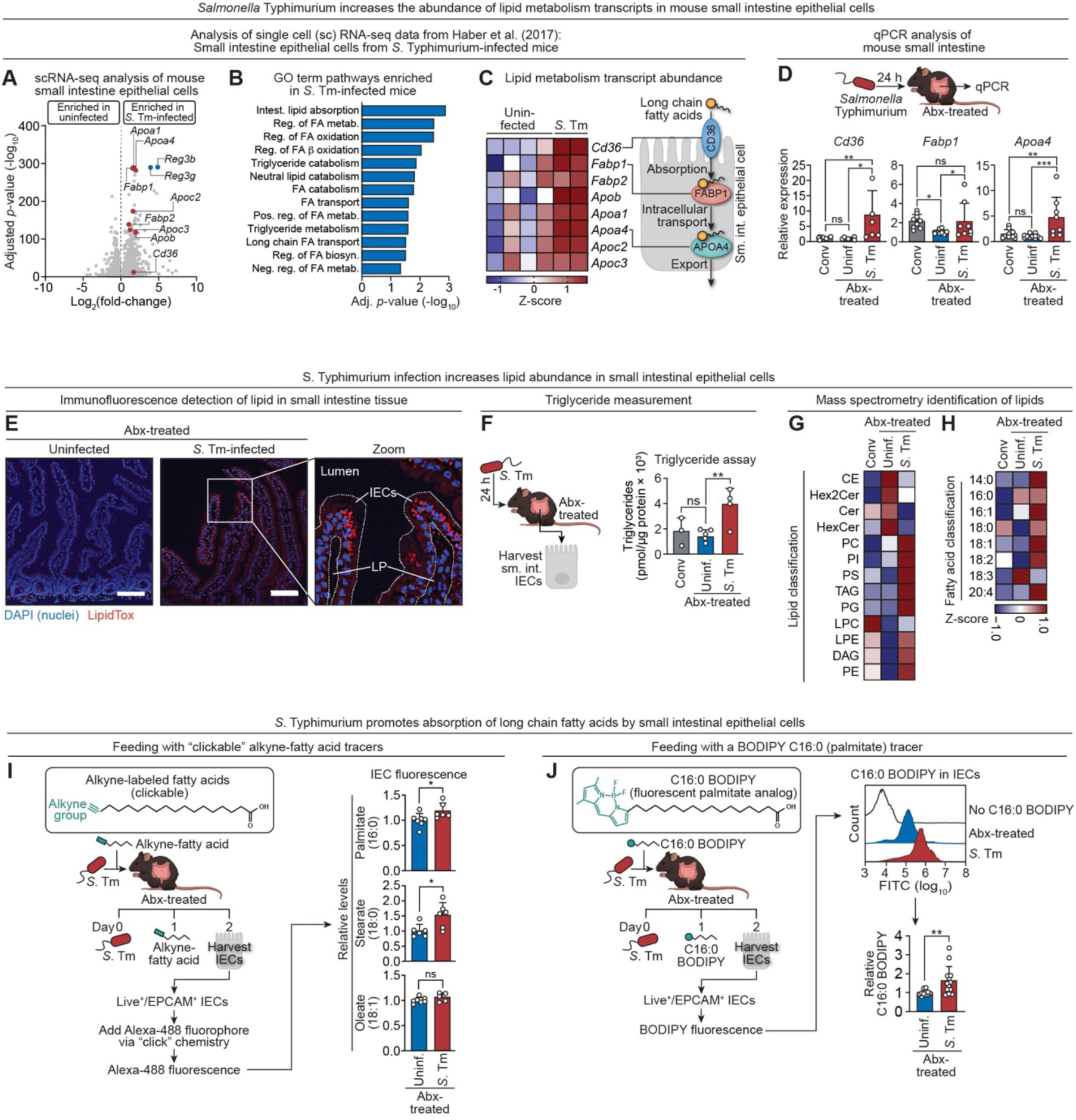
*Salmonella* Typhimurium infection promotes lipid uptake and metabolic reprogramming in small intestinal epithelial cells. **(A)** Volcano plot from re-analysis of published scRNA-seq data of small intestinal epithelial cells (IECs) isolated from streptomycin-pretreated mice 48 h after *S*. Typhimurium SL1344 infection or from uninfected controls (12). Lipid metabolism–associated genes enriched in infected IECs are highlighted in red; *Reg3* genes are shown in blue. **(B)** Gene ontology (GO) pathway analysis of transcripts enriched in IECs from *S*. Typhimurium–infected mice (12). Shown are pathways related to fatty acid metabolism. **(C)** Heatmap showing relative expression (Z-score) of lipid metabolism and transport genes in IECs from uninfected and infected mice (12). Schematic depicts long-chain fatty acid uptake and intracellular trafficking pathways in IECs. **(D)** qPCR analysis of *Cd36*, *Fabp1*, and *Apoa4* expression in small intestine from conventional (Conv) mice or antibiotic-treated (Abx-treated) mice 24 h after infection with *S*. Typhimurium SL1344 (n=5– 12 mice per group). Significance was determined by one-way ANOVA. **(E)** Immunofluorescence microscopy of small intestines from antibiotic-treated uninfected or S. Typhimurium–infected mice, showing neutral lipid accumulation (LipidTOX, red) and nuclei (DAPI, blue). Representative images and zoomed region shown. Scale bar, 100 µm. **(F)** Triglyceride quantification in isolated small intestinal IECs from conventional or antibiotic-treated mice 24 h after *S*. Typhimurium infection (n=3–5 mice per group). Significance was determined by one-way ANOVA. **(G)** Mass spectrometry–based lipidomic analysis of IECs from conventional or antibiotic-treated mice following *S*. Typhimurium infection. Heatmap depicts relative abundance (Z-score) of lipid classes, averaged across 3–4 mice per group. Lipid classes and their abbreviations are defined in Table S1. **(H)** Fatty acid composition of glycerolipids was determined by mass spectrometry for the same IEC samples shown in (G). Heatmap depicts relative abundance (Z-score) of individual fatty acids, averaged across 3–5 mice per group. **(I)** *In vivo* uptake of alkyne-labeled long-chain fatty acids in IECs from antibiotic-treated mice infected with *S.* Typhimurium SL1344. Alkyne-labeled fatty acids were administered by intragastric gavage, followed by click-chemistry detection after IEC isolation. Fluorescence was quantified by flow cytometry as median fluorescence intensity (MFI; n=5–7 mice per group). Significance was determined by Student’s *t*-test. **(J)** *In vivo* uptake of the fluorescent palmitate analog C16:0 BODIPY in IECs from antibiotic-treated mice infected with *S.* Typhimurium SL1344. Mice received C16:0 BODIPY by intragastric gavage, and fluorescence was quantified by flow cytometry (MFI; n=12–13 mice per group). Representative histograms and quantification are shown. Significance was determined by Student’s *t*-test. scRNA-seq, single-cell RNA sequencing; *S*. Tm, *Salmonella* Typhimurium; GO, gene ontology; Sm. int., small intestine; qPCR, quantitative real-time PCR; Abx-treated, antibiotic-treated; Conv, conventional; Uninf, uninfected; IEC, intestinal epithelial cell; LP, lamina propria. Data shown in panels D–F, I, and J are representative of 2–3 independent experiments. Each bar graph data point represents one mouse.

To independently validate these findings while minimizing infection-associated tissue pathology, we analyzed intestinal gene expression early during infection. Mice were treated with antibiotics to deplete the microbiota, were orally infected with *S*. Typhimurium, and intestinal gene expression was measured 24 hours after infection. At this time point, infected mice showed minimal intestinal pathology and similar numbers of viable IECs compared with uninfected controls, indicating that infection did not cause substantial IEC loss (Fig. S1A–D). Nevertheless, intestinal *Il22* expression was robustly induced (Fig. S1E), consistent with the established early interleukin 22 (IL-22) response to *S*. Typhimurium infection and indicating productive host recognition of the pathogen (13).

Infected mice exhibited increased expression of genes involved in lipid transport, including *Cd36*, which mediates long-chain fatty acid uptake (14); *Fabp1*, which facilitates intracellular lipid trafficking (15); and *Apoa4*, which enables lipid export from enterocytes (16) (Fig. 1D). Similar results were observed in germ-free mice, indicating that these transcriptional changes occur independently of the resident microbiota (Fig. S2A). Conversely, conventional mice that were not infected with *S*. Typhimurium did not show increased expression of these genes, indicating that the microbiota alone is insufficient to drive the response (Fig. 1D). Together, these findings indicate that *S*. Typhimurium infection induces a transcriptional program consistent with enhanced lipid uptake and trafficking in intestinal epithelial cells.

The increased expression of *Cd36*, encoding a long-chain fatty acid transporter, suggested that infection might enhance lipid uptake by IECs. To determine whether lipid abundance was altered during infection, we first visualized neutral lipids in intestinal tissue using LipidTOX staining. IECs from infected mice showed marked accumulation of neutral lipids compared with uninfected controls (Fig. 1E). Consistent with this observation, direct biochemical measurement revealed increased accumulation of neutral lipid triacylglycerols (TAGs)—the primary storage form of absorbed dietary fatty acids—in IECs from *S*. Typhimurium–infected mice (Fig. 1F). Unbiased mass spectrometry–based lipidomic analysis further confirmed increased abundance of several lipid classes following infection, including TAGs (Fig. 1G). Infection was also associated with increased abundance of multiple long-chain fatty acid species (Fig. 1H), consistent with increased uptake of dietary fatty acids that are subsequently incorporated into intracellular TAG pools. Together, these data indicate that *S*. Typhimurium infection promotes lipid accumulation in IECs.

Increased intracellular lipid levels can arise from either enhanced uptake of dietary lipids or increased endogenous fatty acid synthesis. To distinguish between these possibilities, we examined expression of genes encoding key enzymes involved in de novo lipogenesis. These included *Fasn*, encoding fatty acid synthase, which synthesizes palmitate (17); *Scd1*, encoding stearoyl-CoA desaturase-1, which generates monounsaturated fatty acids required for cellular lipid biosynthesis (18); and members of the *Elovl* gene family, which elongate fatty acyl-CoAs, producing very-long-chain fatty acids (19). Levels of these transcripts were not appreciably altered in *S*. Typhimurium–infected mice (Fig. S2B), arguing that increased de novo lipogenesis is unlikely to account for the observed lipid accumulation and suggesting that infection instead enhances uptake of dietary fatty acids.

We next tested directly whether *S. Typhimurium* infection modulates the uptake of dietary fatty acids. We focused on three long-chain fatty acids—palmitate (16:0), stearate (18:0), and oleate (18:1)— that were elevated in IECs during infection (Fig. 1H). To quantify uptake, we administered alkyne-labeled fatty acid analogs by intragastric gavage and measured IEC fluorescence by flow cytometry following click-chemistry labeling (Fig. 1I). IECs from infected mice exhibited increased uptake of alkyne-palmitate and alkyne-stearate, whereas alkyne-oleate uptake was unchanged (Fig. 1I). These findings indicate that *S*. Typhimurium infection selectively enhances uptake of specific dietary long-chain fatty acids by the intestinal epithelium.

Because palmitate is a well-characterized substrate for CD36, which preferentially mediates uptake of long-chain fatty acids (14), we next asked whether infection-induced uptake exhibits chain-length specificity. Antibiotic-treated mice received the fluorescent fatty acid analogs C16:0 BODIPY or C12:0 BODIPY by intragastric gavage, and IEC fluorescence was quantified by flow cytometry. *S*. Typhimurium infection increased uptake of C16:0 BODIPY but not C12:0 BODIPY (Fig. 1J; Fig. S2C). C16:0 BODIPY fluorescence remained elevated in IECs from infected mice over eight hours, although signal gradually declined over time, consistent with enterocyte lipid processing and chylomicron-mediated lipid export (20) (Fig. S2D). Together, these findings indicate that *S.* Typhimurium infection selectively enhances epithelial uptake of dietary long-chain fatty acids in a manner consistent with CD36-mediated transport.

### *S*. Typhimurium lipopolysaccharide promotes epithelial long-chain fatty acid uptake

To identify bacterial signals responsible for inducing epithelial cell lipid uptake and metabolism, we first examined whether *Salmonella* virulence mechanisms were required for this response. *S*. Typhimurium invades epithelial cells and survives intracellularly through *Salmonella* pathogenicity island (SPI)-encoded type III secretion systems (21). However, both heat-killed bacteria and a Δ*invA*Δ*spiB* mutant (22) lacking functional type III secretion retained the ability to induce lipid metabolism genes (Fig. S3A–D). These results suggest that bacterial invasion and intracellular survival mechanisms are not required for epithelial lipid metabolic responses.

To determine whether defined bacterial components could recapitulate this phenotype, we tested representative pathogen-associated molecular patterns (PAMPs). These PAMPs included lipopolysaccharide (LPS), a Gram-negative outer membrane glycolipid; flagellin, the protein subunit of bacterial flagella; and lipoteichoic acid (LTA), a Gram-positive cell wall component. Antibiotic-treated mice were injected intraperitoneally with ultra-purified LPS, flagellin, or LTA, and intestinal gene expression and lipid uptake were assessed. Whereas intragastric administration of LPS failed to induce lipid metabolism genes in the small intestine (Fig. S4A), systemic administration of LPS induced expression of lipid transport genes and increased C16:0 BODIPY uptake by IECs (Fig. 2A, B). By contrast, systemic administration of flagellin or LTA had no effect. Consistent with the differential effects of oral and systemic LPS on lipid metabolism genes, oral administration of LPS elicited only a modest increase in intestinal *Il22* expression, whereas systemic administration induced a robust *Il22* response (Fig. S4B). Thus, LPS is sufficient to promote epithelial lipid metabolic gene expression and fatty acid uptake.

**Figure 2.**
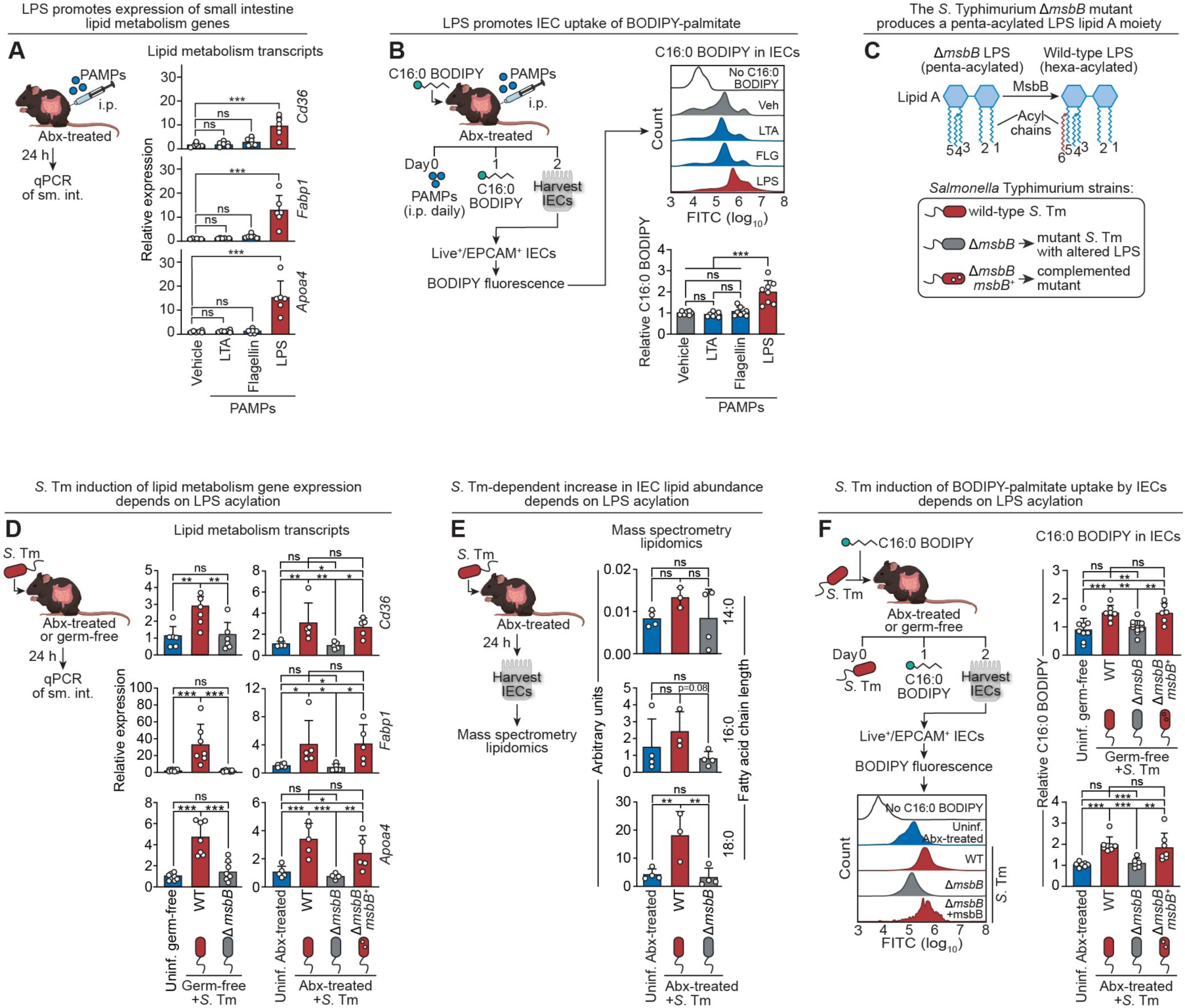
*S*. Typhimurium lipopolysaccharide promotes epithelial long-chain fatty acid uptake. **(A)** qPCR analysis of lipid metabolism mRNAs (*Cd36*, *Fabp1*, and *Apoa4*) in small intestines of antibiotic-treated mice injected intraperitoneally with 1 µg of pathogen-associated molecular patterns (PAMPs): lipoteichoic acid (LTA), flagellin, or ultra-pure lipopolysaccharide (LPS) (n=6 mice per group). **(B)** In vivo uptake of C16:0 BODIPY in small intestinal epithelial cells (IECs) from antibiotic-treated mice injected intraperitoneally with the PAMPs shown in (A). Mice were injected daily for 3 days. Mice were gavaged with C16:0 BODIPY 12 h after the final treatment, and IEC fluorescence was quantified by flow cytometry. Analysis was performed on live EpCAM⁺ cells. Representative histograms and quantification of MFI are shown (n=7–8 mice per group). **(C)** Schematic of *S*. Typhimurium lipopolysaccharide (LPS) biosynthesis and mutant strains used in (D– F). The LPS biosynthesis pathway generates penta-acylated LPS, which is converted to hexa-acylated LPS by the acyltransferase MsbB (LpxM); the Δ*msbB* mutant retains penta-acylated LPS. **(D)** qPCR analysis of lipid metabolism mRNAs (*Cd36*, *Fabp1*, and *Apoa4*) in small intestines of germ-free or antibiotic-treated mice infected by intragastric gavage with parental wild-type or mutant *S*. Typhimurium IR715 strains shown in (C) (n=5–8 mice per group). **(E)** Mass spectrometry–based lipidomic analysis of IECs from uninfected or wild-type or mutant *S*. Typhimurium IR715–infected mice is shown as bar plots. Relative abundance of 14:0, 16:0, and 18:0 fatty acid–containing glycerolipids is shown (n = 3–4 mice per group). **(F)** In vivo uptake of C16:0 BODIPY. Antibiotic-treated mice were infected with parental wild-type or mutant *S*. Typhimurium IR715 strains and orally gavaged with C16:0 BODIPY. IEC fluorescence was quantified by flow cytometry. Representative histograms and MFI quantification are shown (n=6–11 mice per group). LPS, lipopolysaccharide; PAMPs, pathogen-associated molecular patterns; i.p., intraperitoneal; Abx-treated, antibiotic-treated; qPCR, quantitative real-time PCR; sm. int., small intestine; LTA, lipoteichoic acid; IEC, intestinal epithelial cell; Veh, vehicle; *S*. Tm, *Salmonella* Typhimurium; Uninf., uninfected; WT, wild-type. MFI, median fluorescence intensity. Data shown in panels A, B, D, and F are representative of 2–3 independent experiments. Each bar graph data point represents one mouse. Significance was determined by one-way ANOVA.

We next asked whether LPS promotes lipid uptake during live *S*. Typhimurium infection. LPS is essential for bacterial viability, thus precluding generation of a bacterial strain completely lacking LPS. Instead, we used an isogenic *S*. Typhimurium Δ*msbB* mutant, which lacks the LpxM acyltransferase that adds a sixth acyl chain to lipid A (23) (Fig. 2C; Fig. S5A). As a result, the Δ*msbB* mutant produces penta-acylated LPS with reduced innate immune stimulatory activity (24, 25) (Fig. 2C; Fig. S5B, C). Although the Δ*msbB* mutant colonized the intestine at levels comparable to the wild-type and complemented strains (Fig. S5D), it failed to induce lipid transport genes, increase long-chain fatty acid abundance in IECs, or enhance uptake of C16:0 BODIPY (Fig. 2D–F). Thus, *S*. Typhimurium LPS promotes IEC lipid metabolism and fatty acid uptake during infection in an acylation-dependent manner.

### *S*. Typhimurium-induced epithelial lipid uptake requires TLR4 signaling in dendritic cells

We next sought to identify the host sensing pathways that mediate lipid uptake in response to *S*. Typhimurium LPS. LPS can be detected by multiple host pathways. These include TLR4, which signals through the adaptor MYD88 (26, 27), and cytosolic sensing pathways that activate Caspase-11 and downstream inflammasome signaling, leading to maturation and release of IL-1 family cytokines (28, 29). Because both TLR and IL-1 receptor family signaling converge on MYD88, we first asked whether the lipid metabolic response to *S*. Typhimurium requires MYD88-dependent signaling and, if so, whether this reflects TLR-mediated sensing or inflammasome-driven cytokine signaling.

To address this question, we infected *Myd88*^-/-^ mice and mice lacking functional Caspase-11 (129S1) with wild-type *S*. Typhimurium and assessed intestinal lipid metabolism. Mice lacking MYD88 exhibited reduced expression of lipid metabolism genes and decreased uptake of C16:0 BODIPY compared to wild-type controls (Fig. 3A–C), indicating that this response requires MYD88-dependent signaling. In contrast, mice lacking functional Caspase-11 showed no defect in lipid metabolic gene expression or C16:0 BODIPY uptake, demonstrating that Caspase-11–dependent inflammasome activation is dispensable for this phenotype (Fig. S6A–C). These findings indicate that the MYD88 dependence of the response reflects TLR signaling rather than inflammasome-driven IL-1 family signaling.

**Figure 3.**
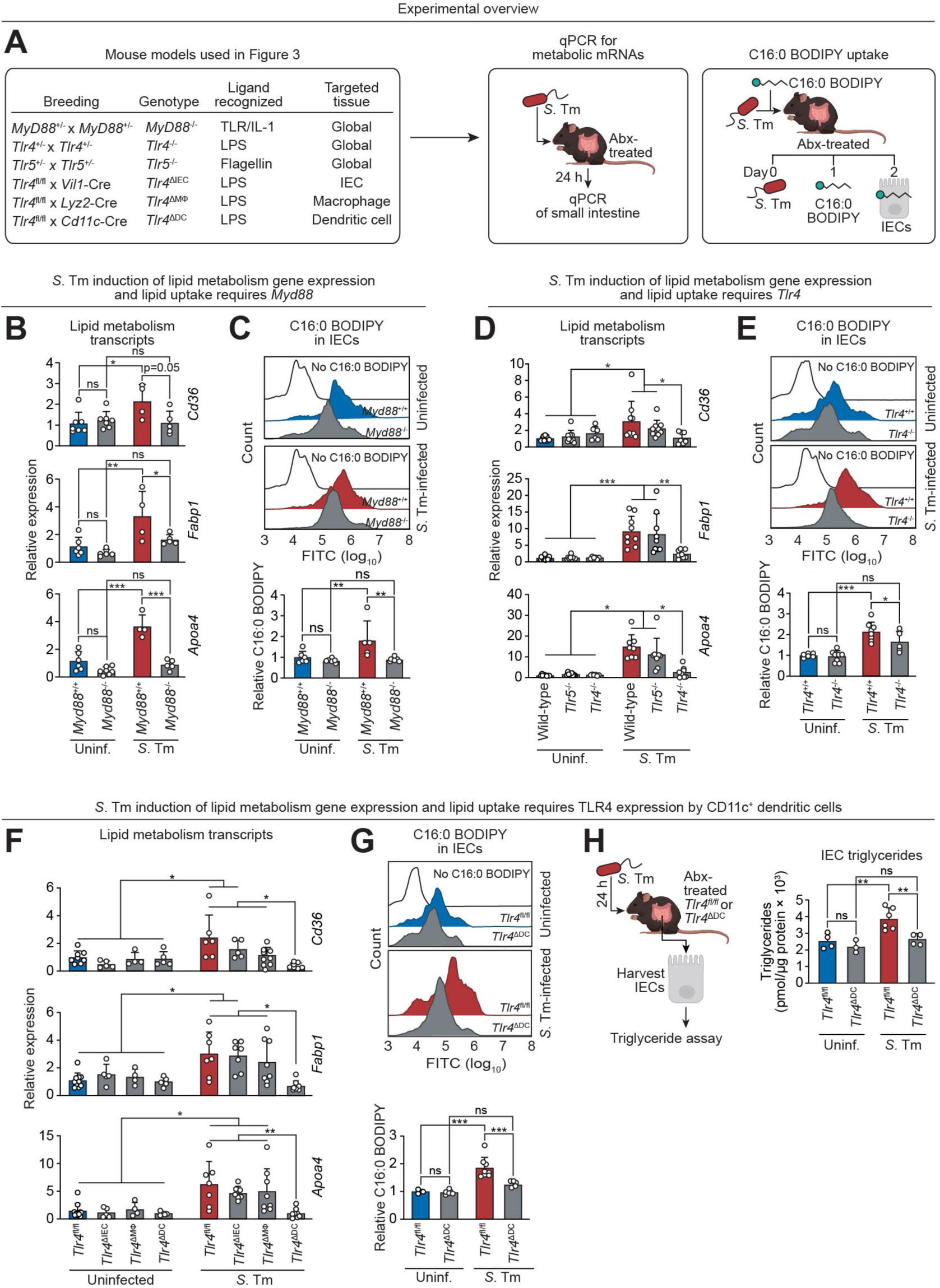
*S*. Typhimurium-induced epithelial lipid uptake requires TLR4 signaling in dendritic cells. **(A)** Mouse models and experimental overview for panels B–H. **(B)** qPCR analysis of lipid metabolism transcripts (*Cd36*, *Fabp1*, and *Apoa4*) in small intestines of antibiotic-treated *Myd88*^+/+^ or *Myd88*^-/-^ littermates 24 h after oral *S*. Typhimurium SL1344 infection (n=4–7 mice per group). **(C)** C16:0 BODIPY uptake by IECs from antibiotic-treated *Myd88*^+/+^ or *Myd88*^-/-^ littermates 24 h after *S*. Typhimurium SL1344 infection. MFI was quantified by flow cytometry; representative histograms and quantification are shown (n=5–6 mice per group). **(D)** qPCR analysis of lipid metabolism transcripts in small intestines of antibiotic-treated wild-type, *Tlr4*^-/-^, or *Tlr5*^-/-^ mice 24 h after *S*. Typhimurium SL1344 infection (n=6–9 mice per group). **(E)** C16:0 BODIPY uptake by IECs from antibiotic-treated wild-type, *Tlr4*^-/-^, or *Tlr5*^-/-^ mice 24 h after *S*. Typhimurium SL1344 infection. MFI was quantified by flow cytometry; representative histograms and quantification are shown (n=5–8 mice per group). **(F)** qPCR analysis of lipid metabolism transcripts in small intestines of antibiotic-treated *Tlr4*^fl/fl^, *Tlr4*^ΔIEC^, *Tlr4*^ΔMφ^, or *Tlr4*^ΔDC^ mice 24 h after *S*. Typhimurium SL1344 infection (n=4–10 mice per group). Only statistically significant comparisons are shown; all other within-genotype comparisons were not statistically significant. **(G)** C16:0 BODIPY uptake by IECs from antibiotic-treated *Tlr4*^fl/fl^, *Tlr4*^ΔIEC^, *Tlr4*^ΔMφ^, or *Tlr4*^ΔDC^ mice 24 h after *S*. Typhimurium SL1344 infection. Fluorescence was quantified by flow cytometry; representative histograms and MFI quantification are shown (n=3–8 mice per group). **(H)** Triglyceride quantification in small intestinal IECs isolated from antibiotic-treated *Tlr4*^fl/fl^ or *Tlr4*^ΔDC^ mice 24 h after *S*. Typhimurium SL1344 infection (n=3–6 mice per group). LPS, lipopolysaccharide; qPCR, quantitative real-time PCR; *S*. Tm, *Salmonella* Typhimurium; Abx-treated, antibiotic-treated; IEC, intestinal epithelial cell; uninf., uninfected; MFI, median fluorescence intensity. Data shown are representative of 2–3 independent experiments. Each bar graph data point represents one mouse. Significance was determined by two-way ANOVA.

We next investigated the specific TLR responsible for sensing *S*. Typhimurium. Because LPS was sufficient to stimulate epithelial lipid uptake (Fig. 2A, B), and *S*. Typhimurium also expresses flagellin, we tested the roles of the corresponding receptors TLR4 and TLR5 by infecting *Tlr4*^-/-^ and *Tlr5*^-/-^ mice (Fig. 3A). Lipid metabolic gene expression and C16:0 BODIPY uptake were reduced in *Tlr4*^-/-^ mice but not in *Tlr5*^-/-^ mice (Fig. 3D, E), identifying TLR4 as the receptor that mediates this response to *S*. Typhimurium.

TLR4 is expressed by multiple cell types in the intestine, including epithelial cells and various myeloid populations. To identify the relevant cellular compartment, we bred *Tlr4*^fl/fl^ mice to mice harboring cell-specific Cre transgenes to generate mice with selective deletions of *Tlr4* in IECs (*Villin*-Cre; *Tlr4*^ΔIEC^), CD11c^+^ dendritic cells (*Cd11c*-Cre; *Tlr4*^ΔDC^), and macrophages (*Lyz2*-Cre; *Tlr4*^ΔMΦ^) (Fig. 3A). Following infection with wild-type *S*. Typhimurium, only *Tlr4*^ΔDC^ mice exhibited reduced lipid metabolic gene expression and decreased uptake of C16:0 BODIPY, whereas *Tlr4*^ΔIEC^ and *Tlr4*^ΔMΦ^ mice were comparable to *Tlr4*^fl/fl^ controls (Fig. 3F, G). Consistent with these findings, triglyceride accumulation in intestinal epithelial cells was reduced in *Tlr4*^ΔDC^ mice during infection (Fig. 3H). Likewise, induction of the infection-responsive cytokine gene *Il22* was impaired in *S*. Typhimurium-infected *Tlr4*^-/-^ and *Tlr4*^ΔDC^ mice (Fig. S7), indicating that TLR4 signaling in CD11c^+^ cells is also required for the intestinal IL-22 response to *S*. Typhimurium. Together, these results show that *S*. Typhimurium LPS is sensed by TLR4 in CD11c⁺ dendritic cells to drive epithelial lipid uptake and metabolic reprogramming.

### The commensal bacterium *Escherichia coli* promotes epithelial long-chain fatty acid uptake via LPS

We next assessed whether a non-invasive commensal bacterium could similarly promote epithelial lipid uptake. We mono-associated germ-free mice with the probiotic commensal *E. coli* Nissle (30, 31) and assessed intestinal lipid metabolism transcripts and epithelial fatty acid uptake. *E. coli* colonized the intestine at levels comparable to *S*. Typhimurium (Fig. 4A, B). This resulted in increased epithelial lipid metabolism gene expression together with modest enhancement of C16:0 BODIPY uptake relative to germ-free controls (Fig. 4A–D). These findings indicate that commensal *E. coli* can stimulate epithelial long-chain fatty acid uptake, although less robustly than *S*. Typhimurium.

**Figure 4.**
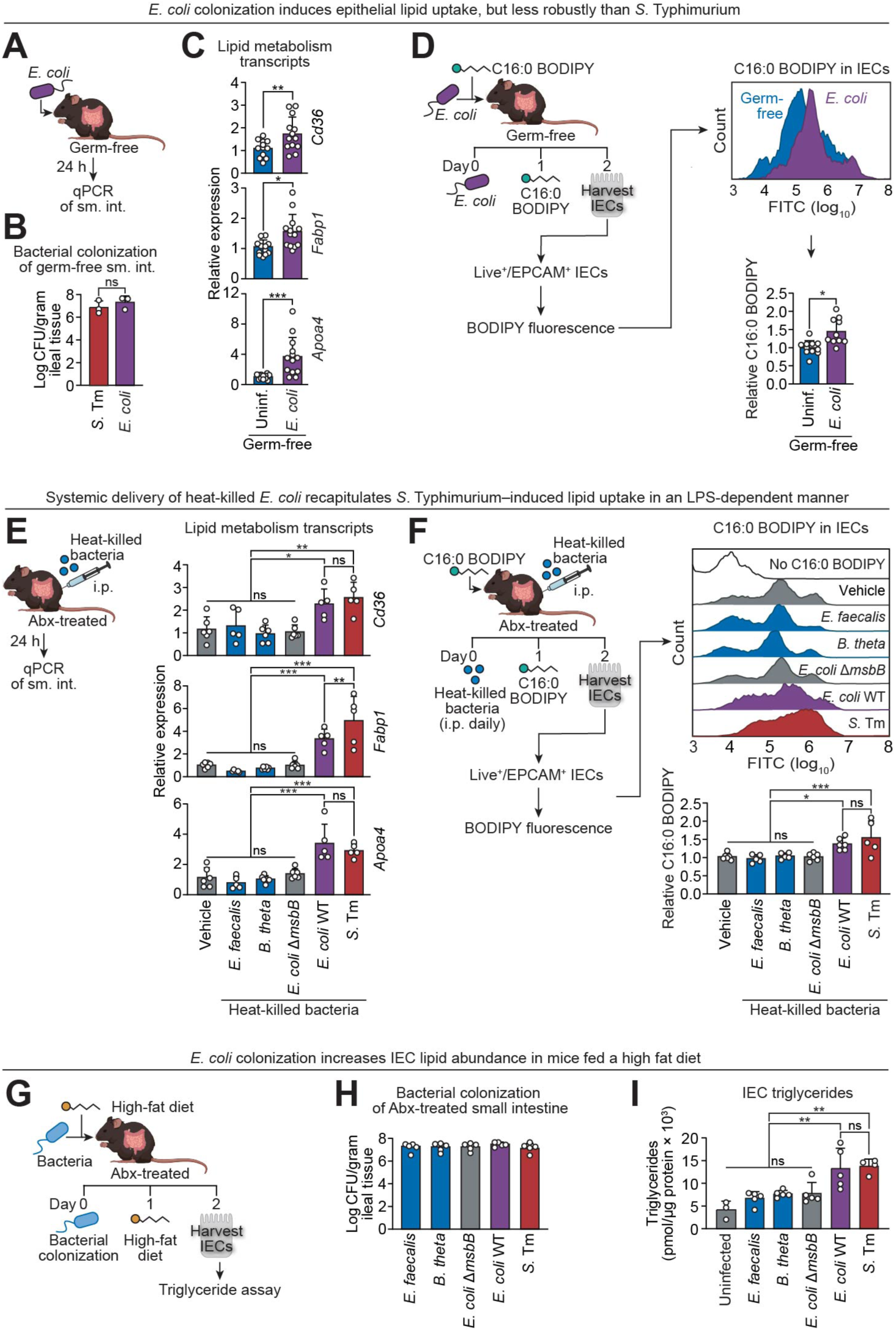
Commensal *Escherichia coli* promotes epithelial long-chain fatty acid uptake via LPS. **(A)** Experimental overview for panels B and C. **(B)** Comparison of bacterial burdens in the small intestines of germ-free mice after oral infection with *S*. Typhimurium SL1344 or *E*. *coli* Nissle. **(C)** qPCR analysis of lipid metabolism transcripts (*Cd36*, *Apoa4*, and *Fabp1*) in small intestines of germ-free or *E. coli* mono-associated mice (n=12–13 mice per group). **(D)** C16:0 BODIPY uptake by IECs from germ-free or *E. coli* mono-associated mice. Mice were gavaged with tracer and IEC fluorescence was quantified by flow cytometry. Representative histograms and quantification of MFI are shown (n=10–11 mice per group). **(E)** qPCR analysis of lipid metabolism transcripts in small intestines of antibiotic-treated mice injected intraperitoneally with 10^7^ CFU of heat-killed bacteria, including *E. coli* (wild-type or Δ*msbB*), *S*. Typhimurium SL1344, *Enterococcus faecalis*, or *Bacteroides thetaiotaomicron* (n=5–6 mice per group). **(F)** C16:0 BODIPY uptake by IECs from antibiotic-treated mice injected with heat-killed bacteria as in (E). Representative histograms and quantification are shown (n=5–6 mice per group). **(G)** Experimental schematic for panels H and I. Antibiotic-treated mice were associated with the indicated bacterial strains and fed a high-fat diet prior to analysis. **(H)** Bacterial burdens in small intestines of mice associated with the indicated bacterial strains (n=5 mice per group). **(I)** Triglyceride abundance in isolated IECs from mice associated with the indicated bacterial strains (n=3–5 mice per group). qPCR, quantitative real-time PCR; sm. int., small intestine; uninf., uninfected; IECs, intestinal epithelial cells; i.p., intraperitoneal; Abx-treated, antibiotic-treated; MFI, median fluorescence intensity. Data shown in panels C–F are representative of 2–3 independent experiments. Each bar graph data point represents one mouse. Significance was determined by Student’s *t*-test for panels B–D and by one-way ANOVA for panels E, F, H and I.

We next asked why *E. coli* induces a weaker lipid metabolic response despite producing structurally similar hexa-acylated LPS (23). Because *S*. Typhimurium invades host tissues whereas *E. coli* remains largely luminal, we hypothesized that *E. coli*-derived LPS has reduced access to the subepithelial myeloid cells that mediate TLR4-dependent signaling. Consistent with this idea, oral administration of purified LPS was insufficient to trigger lipid metabolic responses (Fig. S4A, B), suggesting that luminal exposure alone does not effectively engage this pathway. To test this model directly, we bypassed differences in bacterial localization by systemic administration of heat-killed bacteria. Under these conditions, heat-killed wild-type *E. coli* induced lipid metabolic gene expression and epithelial C16:0 BODIPY uptake to a degree comparable to *S*. Typhimurium (Fig. 4E, F). By contrast, an isogenic *E. coli* Δ*msbB* mutant failed to induce these responses (Fig. 4E, F). Similarly, other commensal species, including the Gram-positive species *Enterococcus faecalis* and *Bacteroides thetaiotaomicron*, which produces weakly immunostimulatory penta-acylated LPS (32, 33), also failed to induce these responses (Fig. 4E, F). Together, these findings suggest that bacterial localization and LPS structure both influence the magnitude of epithelial lipid metabolic responses.

Finally, we addressed whether increased dietary lipid availability amplifies bacterially induced epithelial lipid accumulation. Antibiotic-treated mice were colonized with individual bacterial strains, maintained on a high-fat diet, and IEC triglyceride abundance was measured. Under these conditions, mice colonized with either *E. coli* or *S*. Typhimurium exhibited increased epithelial triglyceride accumulation compared to mice colonized with other commensals or the *E. coli* Δ*msbB* mutant, despite similar intestinal colonization levels (Fig. 4G–I). These findings indicate that commensal bacterial LPS can promote epithelial lipid accumulation, and that the magnitude of this response is shaped by both host exposure and dietary context.

### CD36 contributes to epithelial lipid uptake and limits extraintestinal *S*. Typhimurium burden

Since both *S*. Typhimurium and *E. coli* induce *Cd36* expression in an LPS-dependent manner, we asked whether CD36 functionally contributes to epithelial lipid uptake during infection. To address this, we generated mice with an IEC-specific deletion of *Cd36* (*Cd36*^ΔIEC^) by crossing *Cd36*^fl/fl^ mice with *Villin*-Cre transgenic mice. Under steady-state conditions, IECs from *Cd36*^ΔIEC^ mice exhibited reduced intestinal expression of lipid metabolism genes, including *Fabp1* and *Apoa4* (Fig. 5A), indicating that CD36 contributes to basal epithelial lipid metabolic programming.

**Figure 5.**
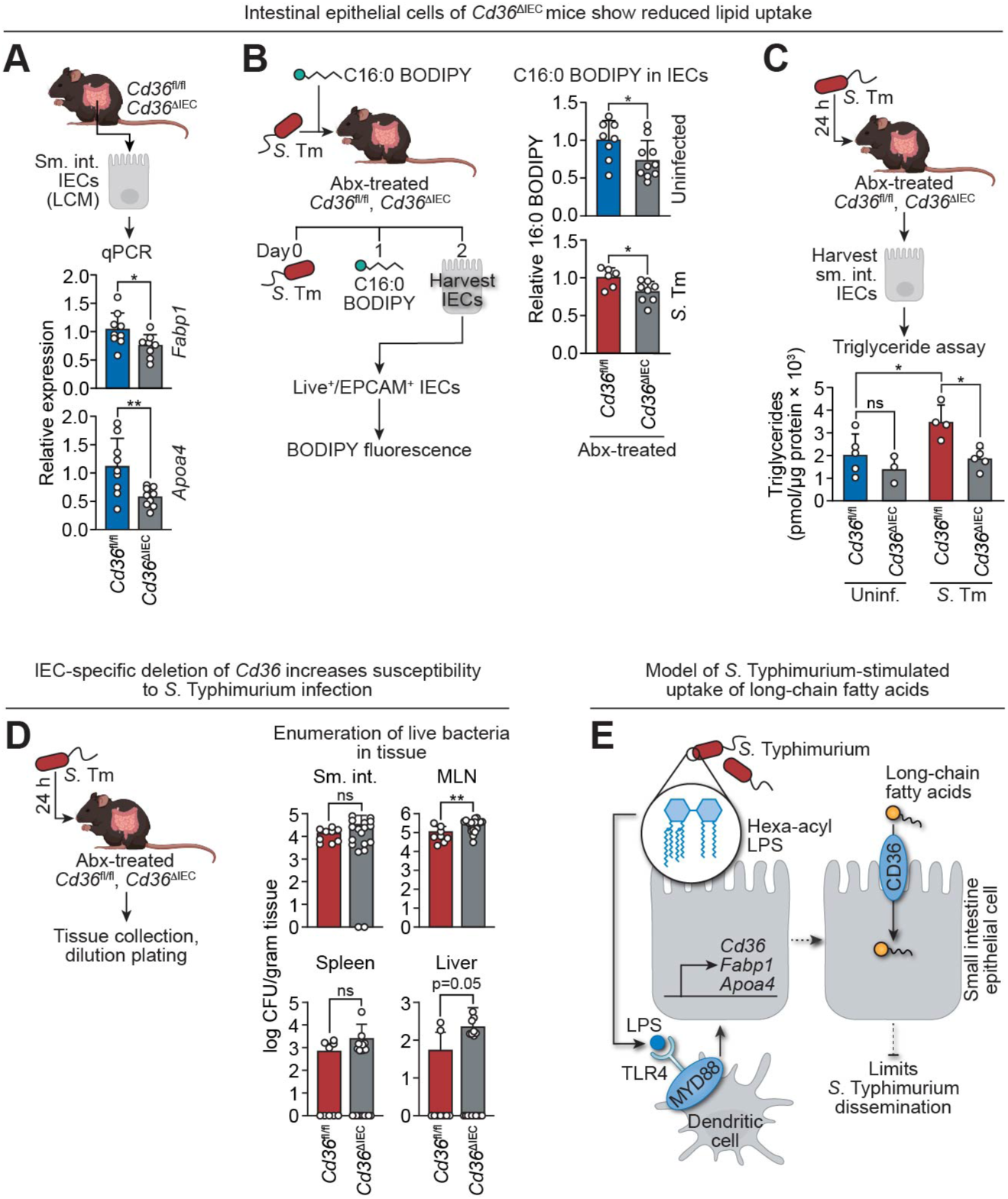
CD36 contributes to epithelial lipid uptake and limits extraintestinal *S*. Typhimurium burden. **(A)** qPCR analysis of lipid metabolism genes (*Apoa4* and *Fabp1*) in IECs recovered by laser capture microdissection from *Cd36*^ΔIEC^ and *Cd36*^fl/fl^ littermates (n=8–9 mice per group). **(B)** In vivo uptake of C16:0 BODIPY by IECs from antibiotic-treated *Cd36*^ΔIEC^ and *Cd36*^fl/fl^ littermates 24 h after *S*. Typhimurium infection. Mice were gavaged with tracer, and IEC fluorescence was quantified by flow cytometry. Representative histograms and MFI quantification are shown (n=6–10 mice per group). **(C)** Triglyceride abundance in IECs isolated from antibiotic-treated *Cd36*^ΔIEC^ and *Cd36*^fl/fl^ littermates 24 h after *S*. Typhimurium infection (n=3–5 mice per group). **(D)** *S*. Typhimurium SL1344 burdens in the small intestine lumen (sm. int.), mesenteric lymph nodes (MLN), spleen and liver 24 h after oral infection of antibiotic-treated *Cd36*^ΔIEC^ and *Cd36*^fl/fl^ littermates (n=8–20 mice per group). **(E)** Model: *S*. Typhimurium–derived hexa-acylated LPS activates dendritic cell TLR4 to induce epithelial lipid uptake programs, promoting long-chain fatty acid uptake. CD36-dependent lipid uptake limits extraintestinal dissemination through an as-yet undefined mechanism. Sm. int., small intestine; IECs, intestinal epithelial cells; LCM, laser capture microdissection; *S*. Tm, *Salmonella* Typhimurium; Abx-treated, antibiotic-treated; Uninf., uninfected; CFU, colony forming units; MFI, median fluorescence intensity. Data shown are representative of 2–3 independent experiments. Each bar graph data point represents one mouse. Significance was determined by Student’s *t-*test for A, B and D, and by two-way ANOVA for C.

We next assessed fatty acid uptake during *S*. Typhimurium infection. *Cd36*^ΔIEC^ mice showed reduced C16:0 BODIPY uptake in IECs under both uninfected conditions and following *S*. Typhimurium infection (Fig. 5B), indicating that CD36 is required for efficient epithelial long-chain fatty acid uptake. Consistent with this finding, *S*. Typhimurium-induced accumulation of triglycerides in IECs was diminished in *Cd36*^ΔIEC^ mice compared to *Cd36*^fl/fl^ controls (Fig. 5C). Together, these findings establish CD36 as a key mediator of epithelial lipid uptake and accumulation during infection.

We next addressed whether CD36-dependent lipid uptake influences host control of *S*. Typhimurium infection. Following oral infection, bacterial burdens in the distal small intestine were comparable between *Cd36*^ΔIEC^ mice and *Cd36*^fl/fl^ littermate controls. However, *Cd36*^ΔIEC^ mice exhibited increased bacterial burdens in the mesenteric lymph nodes and a trend toward higher bacterial numbers in the liver (Fig. 5D), suggesting enhanced extraintestinal dissemination. By contrast, systemic bacterial burdens were unchanged following intravenous infection, which bypasses the intestinal epithelium (Fig. S8). These findings indicate that epithelial CD36 contributes to host defense during *S*. Typhimurium infection and support a model in which bacteria-induced lipid uptake helps to limit dissemination (Fig. 5E).

## Discussion

In this study, we identify a mechanism by which an enteric bacterial pathogen acutely reprograms host intestinal lipid metabolism. We show that infection with *S*. Typhimurium induces rapid uptake of dietary lipids by intestinal epithelial cells. Lipid uptake is driven by lipopolysaccharide (LPS) and depends on its acylation state. This response requires sensing of LPS through the TLR4–MYD88 pathway in intestinal dendritic cells and is accompanied by coordinated upregulation of epithelial lipid metabolism genes, including the long-chain fatty acid transporter CD36, which functionally contributes to limiting bacterial dissemination. These results establish that LPS is a potent regulator of epithelial lipid absorption and reveal a link between innate immune sensing and host nutrient uptake during infection.

These findings expand our understanding of how microbes shape host metabolism by showing that bacterial regulation of lipid absorption depends on the context of host exposure. Previous work revealed that the commensal microbiota promotes lipid uptake through chronic interactions with the host, particularly in the setting of a high-fat diet (5–7). In contrast, our results show that acute exposure to *S*. Typhimurium rapidly enhances epithelial lipid uptake under basal dietary conditions. The observation that commensal *E. coli* induces a similar but attenuated effect suggests that shared microbial signals engage common host pathways, while differences in host exposure determine the magnitude of the response.

Our data suggest that subepithelial sensing of LPS is a key determinant of this difference. Whereas luminal LPS exposure was insufficient to induce epithelial lipid uptake, bacterial invasion or systemic delivery allowed LPS to access subepithelial dendritic cells and stimulate this response. Consistent with this interpretation, luminal LPS elicited only a modest increase in *Il22* expression, whereas systemic administration induced robust *Il22* expression together with epithelial lipid uptake. These results suggest that spatial compartmentalization of microbial sensing is a critical determinant of metabolic outcomes, with invasive pathogens uniquely positioned to activate this pathway by breaching the epithelial barrier.

Our findings further suggest that this pathway may overlap with a previously described microbiota-responsive circuit. In earlier work, we showed that microbiota-induced epithelial lipid metabolism requires IL-22 signaling. Consistent with this model, *Il22* induction during *S*. Typhimurium infection was impaired in both global and CD11c^+^ cell-specific *Tlr4*-deficient mice. Although further studies are needed to determine whether IL-22 is the downstream mediator of *S*. Typhimurium-driven lipid absorption, these findings suggest that pathogen-derived LPS engages components of the same immune signaling circuit.

A central question raised by our findings is the functional significance of increased epithelial lipid absorption during infection. Lipids can fuel bioenergetic demands, remodel membranes to enhance resilience (34), and act as signaling molecules (35). Consistent with a functional role for this pathway, loss of CD36 increased bacterial dissemination during infection. How CD36-dependent lipid uptake contributes to host defense remains unclear but may involve effects on epithelial barrier function, immune cell function, or the availability and fate of bioactive lipid species. An unexpected feature of the lipid uptake response was its selectivity for specific long-chain fatty acids. *S*. Typhimurium infection preferentially enhanced uptake of saturated long-chain fatty acids such as palmitate and stearate, whereas uptake of oleate, a monounsaturated long-chain fatty acid, was not altered under the conditions tested. This substrate selectivity may reflect differences in transporter preference, intracellular trafficking, or metabolic utilization. Whether infection also affects uptake of very long-chain fatty acids remains to be determined.

More broadly, our findings suggest that Gram-negative bacteria with immunostimulatory LPS, particularly members of the Enterobacteriaceae, may be uniquely positioned to influence host lipid metabolism. Expansion of Enterobacteriaceae is associated with metabolic disorders, including obesity, and with conditions characterized by increased intestinal permeability (36–38). Prior work from our group revealed that the microbiota promotes intestinal epithelial lipid accumulation and CD36 expression during high-fat feeding (5–7), suggesting that these observations reflect related mechanisms. Consistent with this idea, repeated intragastric exposure to heat-killed *S*. Typhimurium modestly enhanced weight gain in mice maintained on a high-fat diet (Fig. S9). Although the underlying mechanisms remain undefined, these results suggest that chronic exposure to immunostimulatory bacterial products may contribute to microbiota-associated metabolic phenotypes. Similarly, environmental factors such as dietary emulsifiers, which increase microbial encroachment and low-grade inflammation (39), may enhance host exposure to bacterial LPS and engage related pathways. Together, these findings suggest that bacteria-driven lipid absorption may link intestinal innate immune sensing to metabolic disease.

## Materials and Methods

Additional methods are presented in Supporting Information, Extended Materials and Methods.

### Mice

C57BL/6J, 129SV1, *Myd88*^-/-^, *Tlr4*^-/-^, *Tlr5*^-/-^, *Tlr4*^fl/fl^, *Tlr4*^ΔIEC^, *Tlr4*^ΔDC^, *Tlr4*^ΔMΦ^, *Cd36*^fl/fl^, *Cd36*^ΔIEC^, *Villin*-Cre, *Lyz2*-Cre, and *Cd11c*-Cre mice were bred and maintained under specific pathogen–free (SPF) conditions at the University of Texas Southwestern Medical Center. *Tlr4*^-/-^, *Tlr4*^fl/fl^ (40), *Tlr5*^-/-^ (11, 41), *Cd36*^fl/fl^ (42), *Villin*-Cre (43), *Lyz2*-Cre (LysM-Cre) (44), and *Cd11c*-Cre (45) mice were from Jackson Laboratories. All floxed strains were maintained as homozygotes; all Cre driver lines were maintained as heterozygotes. *Tlr4*^ΔIEC^, *Tlr4*^ΔDC^, and *Tlr4*^ΔMΦ^ mice were generated by crossing *Tlr4*^fl/fl^ mice with *Villin*-Cre mice, *Cd11c*-Cre mice, and *Lyz2*-Cre mice respectively. *Cd36*^ΔIEC^ mice were generated by crossing *Cd36*^fl/fl^ mice to *Villin*-Cre mice.

Germ-free C57BL/6J mice were housed and bred in the gnotobiotic mouse facility at the University of Texas Southwestern Medical Center. All experiments used male and female 10-to 18-week–old mice. All mice were given access to chow and water ad libitum and kept in 12-hour day/night cycles corresponding to 6 AM/6 PM. Mice were euthanized by isoflurane overdose, followed by cervical dislocation. All experiments were performed using protocols approved by the Institutional Animal Care and Use Committees (IACUC) of the University of Texas Southwestern Medical Center.

### Mouse diet

Antibiotic-treated and germ-free mice were fed Rodent Diet 2918 (Inotiv) that was vacuum sealed and irradiated. Chow was tested for sterility by bacterial culture testing and fecal pellets from germ-free mice fed 2918 chow were tested for bacterial growth. For high fat diet studies, mice were fed TD.160495 (Inotiv). Mice were given ad libitum access to all chow.

### Antibiotic treatment of mice

Antibiotics were sourced from Research Products International. Antibiotic cocktail water was made by dissolving 1 g/L neomycin (cat. # N20040), 1 g/L metronidazole (cat. #50-213-513), 1 g/L streptomycin (cat. #62000-100) and 0.5 g/L vancomycin (cat. # V06500) in deionized water followed by sterile filtration with a 0.22 µM mesh filter. Mice were given ad libitum access to antibiotic cocktail water for at least seven days and were treated by intragastric gavage with 100 µL of antibiotic cocktail water at the start of antibiotic treatment. Microbiota depletion was verified by aerobic and anaerobic culture of fecal pellets.

### PAMP treatment of mice

Lyophilized PAMPs were reconstituted in sterile, endotoxin-free water and administered via intraperitoneal (i.p.) injection or intragastric (i.g.) gavage. For i.p. administration, mice received 40 ng/g body weight of ultrapure flagellin (InvivoGen, cat. #tlrl-epstfla), lipopolysaccharide (InvivoGen, cat. #tlrl-pb5lps), or lipoteichoic acid (InvivoGen, cat. #tlrl-pslta). For i.g. gavage, mice were treated with 50 µg/g body weight of LPS or LTA, or 1 µg/g body weight of flagellin. In experiments involving lipid administration, lipids were given 12 h after the final PAMP treatment.

### Bacterial strains

Unless noted otherwise, *S*. Typhimurium and *E. coli* strains were cultured aerobically in Lysogeny Broth (LB; Sigma-Aldrich, cat. #L7658) or on LB plates (Sigma-Aldrich, cat. #L7533) at 37°C. Antibiotics were added to LB and LB plates at the following final concentrations: nalidixic acid (Research Products International, cat. #3374-05-8), 50 mg/L; kanamycin (Kan; Sigma-Aldrich, cat. #K1377), 100 mg/L; carbenicillin (Carb; Goldbio cat. #C-103-5), 100 mg/L and tetracycline (Tet;Research Products International, cat. #T17000), 15 mg/L. *Salmonella enterica* serovar Typhimurium strains IR715 and IR715 *msbB*::Kan^R^ (Δ*msbB*), were from previous studies (46, 47). *E. coli* Nissle 1917 (Wild-type strain O6:K5:H1; EcN) was from a previous study (30, 31). *Bacteroides thetaiotaomicron* cultures (strain VPI-5482) were grown in freshly prepared liquid Tryptone–Yeast extract–Glucose medium. *Enterococcus faecalis* cultures (strain V583) were grown in Brain Heart Infusion broth (Sigma-Aldrich, cat. #53286). Construction of mutant bacterial strains is described in SI Appendix, Extended Materials and Methods.

### Treatment of mice with alkyne– and BODIPY–conjugated fatty acids

Lyophilized alkynated lipids were from Avanti Research: oleic acid (17-yne), cat. #900412E; palmitic acid (15-yne), cat. #900400P; stearic acid (17-yne), cat. #700383P). Fatty acids were reconstituted to 1 mg/mL DMSO. Mice were given 800 nM of lipid in a 100 µL bolus by intragastric gavage. Click-chemistry attachment of CalFluor 488 azide (Click Chemistry Tools; cat. #1369-1) to alkyne-lipids was performed with Click-iT Cell Reaction Buffer Kit (Thermo Scientific; cat. #C10269) following the manufacturer’s recommended protocol with the following changes. Briefly, following preparation of IECs for flow cytometry analysis (see below), cells were fixed and permeabilized with 4% PFA and 0.02% saponin for 15 minutes at room temperature, protected from light. Cells were washed once in 1% BSA/PBS solution then incubated with Click-iT Reaction Cocktail containing 50 µM CalFluor 488 azide for one hour at room temperature protected from light. Cells were washed once in 1% BSA/PBS solution then resuspended in FACS buffer (PBS supplemented with 3% FBS and 1 mM EDTA) for flow cytometry analysis.

BODIPY-conjugated fatty acids were from Thermo Scientific: BODIPY™ FL C16 (C16:0 BODIPY; cat. #D3821) or BODIPY™ FL C12 (C12:0 BODIPY; cat. #D3822) were reconstituted in DMSO to 1 mg/ml. Mice were given 4 µg of BODIPY-conjugated fatty acid per gram body mass by intragastric gavage and were sacrificed 4 hours later. Mice that were infected with *S*. Typhimurium and treated with BODIPY-conjugated fatty acids were excluded from analysis if their stomachs showed visible signs of gastroparesis.

### Isolation of intestinal epithelial cells

Mouse small intestines were harvested immediately following euthanasia by excising tissue at the stomach and cecum. Mesenteric adipose tissues were removed and the intestinal lumen was flushed with cold wash buffer (PBS supplemented with 3% fetal bovine serum [FBS]) to remove luminal contents. Intestines were then opened longitudinally, and tissues were washed twice with gentle agitation in a 100 mm dish containing 10 mL of wash buffer.

Tissues were cut into 2–3 cm segments, transferred to 50 mL conical tubes, and washed twice by vortexing in 10 mL of wash buffer. Epithelial cells were subsequently dissociated by incubation in 10 mL of IEC separation buffer (3% FBS and 5 mM EDTA in Hank’s balanced salt solution) with shaking at 37°C (225 RPM, 45° angle) for 20 minutes. The supernatant containing IECs was collected and passed through a 70 µm cell strainer to remove residual tissue. Remaining tissue fragments were washed twice with cold wash buffer to remove residual EDTA and recover additional IECs.

For lipidomics analysis, isolated IECs were pelleted at 4°C, and washed once by resuspension in 1 mL cold PBS. Cell numbers were determined, and aliquots of 1 × 10^6^ cells were pelleted at 4°C and flash-frozen in liquid nitrogen.

### Flow cytometry of intestinal epithelial cells

IECs were isolated as described above, and 1% of the total cell suspension (200 µL of a 20 mL suspension) was used for flow cytometry analysis. Cells were washed once with PBS and stained with a viability dye (Ghost Dye 710, Tonbo Biosciences, cat. #13-0871), followed by F_c_ receptor blockade (anti-CD16/32; BD Bioscience cat. #553142) in PBS. After a 20-minute incubation at room temperature, cells were washed once with PBS and stained with anti-EpCAM (PE/Cyanine-7 anti-mouse CD326, BioLegend, cat. #118216) in FACS buffer for 20 minutes at room temperature. Cells were then washed once in FACS buffer and resuspended in FACS buffer for analysis. Flow cytometry was performed using a NovoCyte instrument and data were analyzed with FlowJo software.

### Triglyceride assay

Intestinal epithelial cells were isolated as described above. Cells were pelleted, and supernatants were removed prior to flash freezing. Cell pellets were processed according to the manufacturer’s protocol using a Triglyceride Assay Kit (Abcam, cat. #ab178780). Triglyceride levels were normalized to total protein concentration as determined by Bradford Assay.

## Acknowledgements

We thank J. Shelton (UT Southwestern Histology Core) for assistance with preparing tissue slides. Funding: This work was supported by NIH grants R01 DK070855 (L.V.H.), R01 DK134692 (W.Z.), R35 GM147470 (W.Z.), and R01 AI118807 (S.E.W.); Welch Foundation Grant I-1874 (L.V.H.); Burroughs Wellcome PATH Award #1017880 (S.E.W.); the Walter M. and Helen D. Bader Center for Research on Arthritis and Autoimmune Diseases (L.V.H.); and the Howard Hughes Medical Institute (L.V.H.). E.K. was supported by NIH F31 DK126391, G.Q. was supported by NIH T32 AI005284, and K.N.K was supported by Crohn’s and Colitis Foundation of America Research Fellowship award #1440322. J.G.M. acknowledges support from P30 DK127984 and P01 HL160487. Components of some figures were generated using BioRender.

