## Supporting Information for "*Salmonella* lipopolysaccharide stimulates uptake of long-chain fatty acids in the small intestine"

**This PDF file includes:**

Supporting text
Figs. S1 to S9
Tables S1 and S2
SI References

### Extended Materials and Methods

#### Construction of mutant bacterial strains

IR715 Δ*msbB* + pEK3 was created in this study. pEK3 was created by Gibson assembly using pWSK29 (1)(Addgene, cat. #172972) as a backbone and a fragment of *msbB* from IR715 containing the gene and 150 nucleotides upstream and downstream of the gene. The wild type, Δ*msbB*, and Δ*msbB* + pEK3 strains were grown overnight from a single colony in 5 mL of LB broth with 100 µg/mL nalidixic acid, 50 µg/mL kanamycin, and/or 100 µg/mL carbenicillin overnight at 37°C in a 225 RPM shaker. Overnight cultures were grown to log phase for 3 hours at 37°C in a shaker at 225 RPM. 1 mL of overnight culture of *S*. Typhimurium Δ*msbB* and 250 µL Δ*msbB* + pEK3 were inoculated into a 25 mL flask of low salt LB broth supplemented with $Mg^{2+}$ and $Ca^{2+}$ for growth to log phase. Log phase bacteria were pelleted, washed once in PBS, and then resuspended in PBS. *S.* Typhimurium strain SL1344 (2) and SL1344 *invA*::Tet$^R$ *spiB*::Kan$^R$ (3) were grown overnight from a single colony in 5 mL of LB broth with 100 µg/mL streptomycin (Research Products International, cat. #62000-100) overnight at 37°C on a shaker at 225 RPM.

*E. coli* Nissle 1917 (Wild-type strain O6:K5:H1; EcN) was from a previous study (4, 5). The *E. coli* Nissle Δ*msbB* mutant was created in this study (Nissle 1917 *msbB*::pSW1435). The upstream and downstream regions of the EcN *msbB* (*lpxM*) gene were amplified by PCR from the EcN genome using the primers listed in Table S2. These DNA fragments were introduced into SphI-digested pGP705 (6) using the Gibson assembly reaction (New England Biolabs). The resulting plasmid, pSW1435, was initially propagated in DH5α λ*pir* (7) and then transferred into S17-1 l*pir* (8) for the subsequent conjugation. pSW1435 was introduced into EcN (pSW172) using conjugation on LB plates incubated at room temperature. Colonies in which pSW1435 had integrated into the *msbB* gene in the EcN chromosome were identified on LB plates supplemented with kanamycin and carbenicillin. The helper plasmid pSW172 was cured from the resulting Nissle 1917 *msbB*::pSW1435 mutant (SW1437) by passaging at 37$^o$C. Insertion of the plasmid was confirmed by PCR.

#### Inoculation of mice with bacteria

Mice were housed in ABSL2 rooms prior to infection. Mice that were on antibiotic water were switched to normal water for 24 hours prior to infection. Mice were given 200 µL of bacterial culture resuspended in phosphate buffered saline (PBS) by oral gavage. Mice treated with *S*. Typhimurium strain SL1344 received $10^5$ CFU by intragastric gavage. Conventional mice treated with *S*. Typhimurium IR715 strains received $10^5$ CFU of wild-type and Δ*msbB* + pEK3 or $10^8$ CFU of Δ*msbB* by intragastric gavage. Gnotobiotic mice that were mono-associated with IR715 strains received $10^3$ CFU of wild-type, Δ*fliC*, and ΔmsbB + pEK3 strains and $10^5$ CFU of the Δ*msbB* strain. Mice infected systemically with *S.* Typhimurium received $10^3$ CFU of wild-type SL1344 strain by tail-vein injection.

#### Bacterial burden assays

Mice infected by intragastric gavage were sacrificed two days post-infection. Tissue was homogenized for 15 or more seconds in 1 mL of sterile PBS and 10 µL of homogenate was spread on appropriate antibiotic containing LB agar plates in duplicate. CFUs were counted the next day and burden was normalized to the mass of the tissue. Mice infected by tail-vein injection were sacrificed one day after infection and all other procedures were the same.

#### Treatment of mice with heat-killed bacteria

Overnight bacterial cultures were pelleted, washed once in PBS, and then resuspended in 1 mL of PBS. Dilution plating was performed to determine CFUs. Bacterial suspensions were then warmed to 90$^o$C on a heat block for 10 minutes. Suspensions were placed on ice for 5 minutes and then placed on heat block for another 10 minutes. After the second heat treatment, suspensions were placed at -80$^o$C. Once CFU counts were determined, heat-killed suspensions were adjusted to have matching CFU concentrations of $10^8$ CFU per mL. Heat-killed suspensions were checked for surviving bacteria by culturing a 10 µL aliquot in both liquid culture and on agar plates. Mice were injected with 100 µL of this suspension resulting in treatment with $10^7$ CFUs of dead bacteria.

#### Mass spectrometry lipidomics

All solvents were LC/MS or HPLC grade and obtained from Sigma-Aldrich (St Louis, MO, USA). Splash® Lipidomix® internal standards were purchased from Avanti Polar Lipids (Alabaster, AL, USA).

Lipid extractions were carried out in 16 × 100 mm glass tubes fitted with PTFE-lined caps (Fisher Scientific, Pittsburgh, PA, USA. For each sample, material equivalent to 250,000 cells was transferred into a fresh glass tube prior to extraction.

Lipids were isolated using a modified methyl tert-butyl ether (mTBE) liquid-liquid extraction protocol (9). Samples were combined with 1 mL of water, 1 mL of methanol, and 2 mL of mTBE, followed by vortexing and centrifugation at 2,671 × g for 5 min to achieve phase separation. The upper organic layer was collected into a new glass tube and supplemented with 20 µL of Splash® Lipidomix® standards diluted 1:5 in solvent. Extracted lipids were evaporated under a nitrogen stream and reconstituted in 400 µl of hexane.

Lipidomic analyses were performed using a SCIEX QTRAP 6500$^+$ mass spectrometer (SCIEX, Framingham, MA) interfaced with a Shimadzu LC-30AD HPLC system (Shimadzu, Columbia, MD). Chromatographic separation was conducted on a Supelco Ascentis silica column (150 × 2.1 mm, 5 µm particle size; Supelco, Bellefonte, PA, USA). Samples were resolved at a flow rate of 0.3 mL/min, with an initial mobile phase composition of 97.5% solvent A (hexane) and 2.5% solvent B (mTBE). The gradient program increased solvent B to 5% over 3 min, followed by a ramp to 60% over the subsequent 6 min. Solvent B was then reduced to 0% within 30 sec while Solvent C (90:10 (v/v) isopropanol-water) was introduced at 20% and increased to 40% during the next 11 min. Solvent C was subsequently raised to 44% over 6 min and then to 60% over 50 sec. The column was maintained at 60% solvent C for 1 min before re-equilibration to starting conditions (2.5% solvent B) for 5 min at a flow rate of 0.6 mL/min. Solvent D [95:5 (v/v) acetonitrile-water containing 10 mM ammonium acetate] was infused post-column at 0.03 ml/min. The column compartment was maintained at 25°C throughout data acquisition.

Mass spectrometry data were collected in both positive and negative ionization modes using multiple reaction monitoring. Lipid species were quantified with MultiQuant software (SCIEX), and signal intensities were normalized to the corresponding internal standard species.

For downstream analyses, mass spectrometry values were normalized to total protein content determined by Bradford assay. Normalized datasets were subsequently centered, scaled, and transformed to Z-scores. Mean Z-scores for each experimental condition were visualized using the pheatmap package in R (https://CRAN.R-project.org/package=pheatmap).

**Reanalysis of small intestinal single-cell RNAseq data from Haber et al. (10)**

Raw count matrices were obtained from the Gene Expression Omnibus (GEO) under accession GSE92332 and Single Cell Portal (SCP) under accession SCP44. All cells included by the original authors that met minimum feature count thresholds were retained for downstream analysis. Data were analyzed using the Seurat v5 R package (11, 12). Gene expression counts for each cell were log scale normalized and multiplied by a scale factor of 10,000. Data were subsequently centered and scaled prior to downstream analyses. Dimension reduction was performed for the entire experiment using principal component analysis (*RunPCA*), followed by uniform manifold approximation and projection (*RunUMAP*).

Differential expression analysis was then performed using the DESeq2 algorithm within *FindMarkers*. Gene set enrichment analysis was performed using the *GSEA* function from the clusterProfiler R package (13, 14) with reference *Mus musculus* Biological Process Gene Ontology C5 gene sets obtained from the Molecular Signature Database (MSigDB)(15) via the msigdbr R package. Visualizations of data were generated within Seurat (11, 12), ggplot2 (16), and pheatmap R packages (https://CRAN.R-project.org/package=pheatmap).

**Laser capture microdissection**

Laser capture microdissection was performed essentially as previously described (17). Approximately 5 cm sections of distal ileum were excised, and the lumen was flushed with cold wash buffer. The ileal segments were filled with optimal cutting temperature (OCT) compound and snap-frozen. Frozen tissues were sectioned at 7 µm using a cryostat and mounted onto Permafrost glass slides. Sections were fixed in 70% ethanol, stained with methyl green and eosin following hydration, and subsequently dehydrated through graded alcohols and air-dried. Laser capture microdissection was performed immediately using the Arcturus PixCell II system. For each sample, approximately 10,000–15,000 infrared laser pulses were used to capture cells onto two caps, which were pooled for downstream analysis. Total RNA was extracted with the PicoPure RNA Isolation Kit following the manufacturer’s protocol.

**RNA extraction, cDNA synthesis, and qPCR**

Mice were euthanized, and the terminal 2 cm of small intestine was excised immediately proximal to the cecum. Luminal contents and mucus were gently expelled, and tissue was placed in RNALater for stabilization. Total RNA was purified from whole ileal tissue using RNeasy Mini Kit (Qiagen, cat. #74116) on a QIAcube Connect automated RNA/DNA extraction instrument (Qiagen). Complementary DNA was synthesized using the M-MLV reverse transcription kit (Thermo Fisher Scientific, cat. #28025-021) following the manufacturer's protocol, using 1 µg of total RNA from whole tissue samples or 100–300 ng of total RNA from laser capture microdissected samples. Quantitative PCR was performed using TaqMan Gene Expression Assays (Applied Biosystems, Thermo Fisher Scientific) for *Gapdh*, *Fabp1*, *Apoa4*, and *Cd36* in combination with TaqMan Universal PCR Master Mix (Thermo Fisher Scientific, cat. #4369542) according to the manufacturer's instructions. Gene expression was normalized to 18S rRNA (*Rn18s*) using the $\Delta\Delta C_t$ method. Reactions were run on a QuantStudio 7 Flex Real-Time PCR System (Applied Biosystems, cat. #4485701).

**Immunofluorescence Microscopy**

Small intestines were excised, and the terminal 10 cm of ileum was isolated. Tissue was cut longitudinally to expose the lumen and washed twice in 10 cm Petri dishes containing 10 mL of cold wash buffer. Residual mucus was gently removed, and tissues were fixed in neutral buffered fixative for 8 hours at 4oC with gentle rocking. Following fixation, tissues were transferred to sterile-filtered 30% sucrose in PBS for cryopreservation. Once equilibrated, samples were embedded in OCT compound and cryosectioned.

Cryosections were brought to room temperature in PBS for 5 minutes prior to staining. For neutral lipid staining, hydrated tissues were incubated with LipidTox (Thermo Scientific, cat. #H34476) (1:200 in PBS) for 30 minutes at room temperature. Slides were washed twice in PBS and mounted with glass coverslips using mounting medium containing DAPI. Imaging was performed using a Keyence BZ-X1000 fluorescence microscope.

**HEK TLR4 reporter assay**

HEK Dual™ mTLR4 reporter cells (InvivoGen, cat. #Hkd-mtlr4ni) were maintained according to the manufacturer's instructions. Cells were cultured in DMEM supplemented 10% FBS and 1% penicillin–streptomycin at 37°C in a humidified 5% $CO_2$ incubator. Following initial recovery and two passages, cells were maintained in DMEM with 10% FBS, 1% penicillin–streptomycin, 10 µg/mL blasticidin, 200 µg/mL hygromycin B gold, 1 µg/mL puromycin, and 100 µg/mL zeomycin. One passage prior to experiments, cells were cultured in antibiotic-free DMEM supplemented with 10% FBS. For assays, cells were seeded into 96-well plates at a density of 20,000 cells per well in DMEM containing 10% FBS. Cells were then treated with bacterial PAMPs or bacterial cultures, as indicated, for 4–6 hours. Following stimulation, cell debris was removed by centrifugation at 100*g* for 1 minute, and supernatants were collected for measurement of secreted alkaline phosphatase activity, which reports on NF-κB activation. Detection was performed according to the manufacturer's protocol.

**Weight gain analysis**

To ensure baseline consistency, mice with starting weights within 5% of each other were randomized into control (PBS) or treatment (heat-killed *S.* Typhimurium SL1344) groups. S. Typhimurium-treated mice were administered $10^{11}$ CFU of heat-killed bacteria. Treatments were administered twice weekly by intragastric gavage, with body mass recorded weekly following the second dose. The treatment continued for seven weeks. Mice were maintained on a high-fat diet for the duration of the experiment.

Data were first assessed for normality using the Shapiro-Wilk test. Upon confirmation of normal distribution, percent body weight change over time was analyzed by two-way ANOVA with Sidak's post-hoc test for multiple comparisons. Absolute body weight change at week 5 was compared between groups using Student's *t*-test.

**Quantification and statistical analysis**

Details of statistical analyses for individual experiments, including definitions of significance and specific tests used, are provided in the corresponding figure legends. Data are presented as mean ± standard error of the mean (SEM). The number of experiments indicated in the figure legends represents independent biological replicates performed on separate days. Statistical significance was defined as follows: $*p < 0.05$; $**p < 0.01$; $***p < 0.001$; ns, not significant.

Sample sizes were not determined using statistical methods. Experiments were not conducted in a blinded manner. While formal randomization procedures were not applied, mice were assigned to experimental groups in a random manner, and sample processing order was not predetermined.

All statistical analyses were performed using GraphPad Prism software (version 7.0). Comparisons between two groups were evaluated using two-tailed Student's *t*-tests. For comparisons involving multiple groups, one-way or two-way analysis of variance (ANOVA) was used as appropriate, followed by Tukey's post hoc multiple comparison test (except in Fig. S9B, where Sidak's post hoc test was used as indicated in the legend). Mice that died during experiments were excluded from analysis.

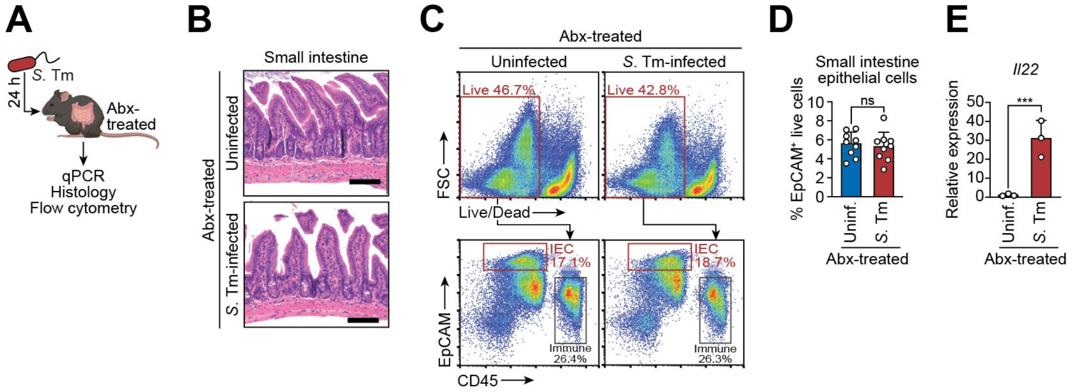


**Fig. S1. *S*. Typhimurium infection does not induce intestinal pathology or epithelial cell death at 24 hours.**

**(A)** Experimental overview for panels B–E. Antibiotic-treated (Abx-treated) mice were orally infected with *S*. Typhimurium (*S*. Tm) for 24 hours. Small intestines were analyzed by qPCR, histology and flow cytometry.

**(B)** Representative histological images of distal small intestines from uninfected and *S*. Typhimurium SL1344-infected mice.

**(C)** Representative flow cytometry plots showing the gating strategy for live EpCAM$^+$ intestinal epithelial cells (IECs) from uninfected and *S*. Typhimurium SL1344-infected mice.

**(D)** Quantification of live EpCAM$^+$ IECs from uninfected (Uninf.) and *S*. Typhimurium SL1344 infected mice (n=9 mice per group). Each bar graph data point represents one mouse. Significance was determined by Student's *t*-test; ns, not significant.

**(E)** qPCR analysis of *Il22* expression in small intestines from uninfected antibiotic-treated (Abx-treated) mice or antibiotic-treated mice 24 h after *S*. Typhimurium SL1344 infection (n=3 mice per group), confirming induction of an early host immune response. Significance was determined by Student's *t*-test; ***, $p < 0.001$.

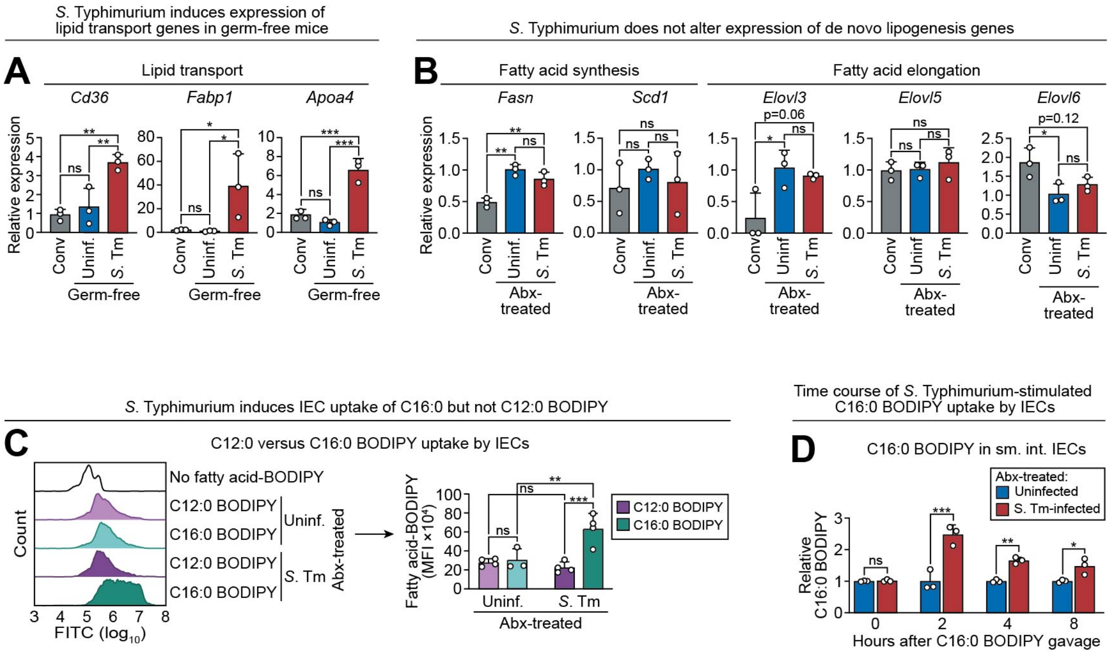


**Fig. S2. Supporting analyses of lipid uptake and metabolism following *S*. Typhimurium infection.**

**(A)** qPCR analysis of genes involved in lipid transport (*Cd36*, *Fabp1*, and *Apoa4*) in small intestines from conventional (Conv) mice, uninfected germ-free mice, or *S*. Typhimurium SL1344-infected germ-free mice (n=3 mice per group). Significance was determined by one-way ANOVA.

**(B)** qPCR analysis of genes involved in de novo lipogenesis (*Scd1*, *Fasn*, *Elovl3*, *5*, and *6*) in small intestines from conventional (Conv) mice, uninfected antibiotic-treated mice, or S. Typhimurium SL1344-infected antibiotic-treated mice (n=3 mice per group). Significance was determined by one-way ANOVA.

**(C)** IEC uptake of C12:0 BODIPY versus C16:0 BODIPY. Antibiotic-treated mice were infected and orally gavaged with BODIPY-conjugated fatty acid tracers; IEC fluorescence was quantified by flow cytometry (n=3–4 mice per group). Representative histograms are shown on the left and quantification on the right. Significance was determined by two-way ANOVA.

**(D)** Antibiotic-treated mice were infected with *S*. Typhimurium SL1344 and orally gavaged with C16:0 BODIPY tracer as shown in Fig. 1J. IEC uptake was measured over 8 hours (n=3 mice per group per time point). Significance was determined by two-way ANOVA.

*S*. Tm, *Salmonella* Typhimurium; Abx-treated, antibiotic-treated; IECs, intestinal epithelial cells; Uninf., uninfected; Conv, conventional. Each bar graph data point represents one mouse.

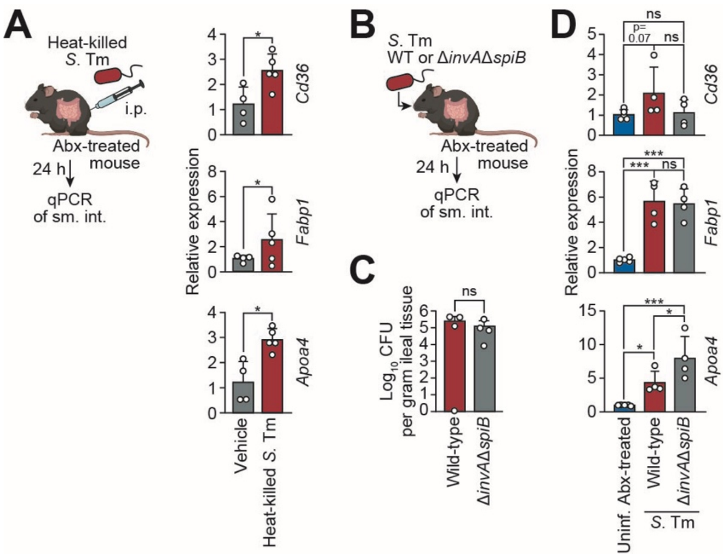


**Fig. S3. *S*. Typhimurium type III secretion is dispensable for lipid metabolic gene induction.**

(A) qPCR analysis of lipid metabolism genes (*Cd36*, *Apoa4*, and *Fabp1*) in distal small intestines from mice treated with heat-killed *S*. Typhimurium SL1344 by intraperitoneal injection. Significance was determined by Student's *t*-test.

(B) Experimental overview for panels (C) and (D). Antibiotic-treated mice were orally infected for 24 h with parental wild-type or ΔinvAΔspiB mutant *S*. Typhimurium SL1344 strains (deficient in type III secretion). Small intestines were analyzed by qPCR. Significance was determined by Student's *t*-test.

(C) Bacterial burden in distal small intestines of mice infected with wild-type or ΔinvAΔspiB *S*. Typhimurium SL1344 for 24 h. Significance was determined by Student's *t*-test.

(D) qPCR analysis of *Cd36*, *Apoa4*, and *Fabp1* in distal small intestines from mice infected with wild-type or ΔinvAΔspiB *S*. Typhimurium SL1344. Significance was determined by one-way ANOVA.

*S*. Tm, *Salmonella* Typhimurium; Abx-treated, antibiotic-treated; qPCR, quantitative real-time PCR; WT, wild-type; CFU, colony forming units; Uninf., uninfected. Each data point represents one mouse.

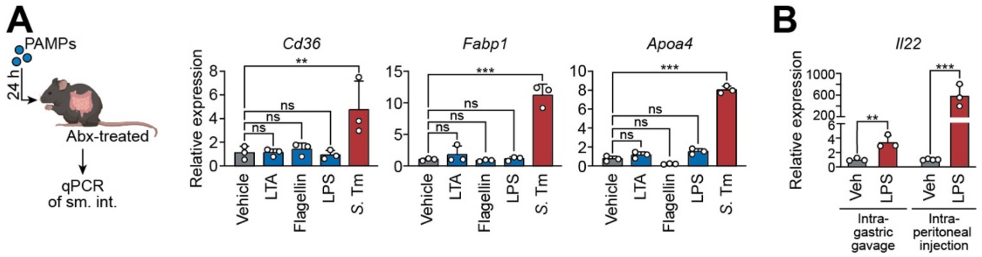


**Fig. S4. Oral PAMP administration fails to recapitulate infection-induced lipid metabolic gene expression.**

(A) qPCR analysis of lipid metabolism genes (*Cd36*, *Apoa4*, and *Fabp1*) in distal small intestines of antibiotic-treated mice gavaged with pathogen-associated molecular patterns: lipoteichoic acid (1 mg), flagellin (20 µg), or ultra-pure lipopolysaccharide (1 mg). Mice infected orally with *S*. Typhimurium as in Fig. 1D are shown for comparison. Significance was determined by one-way ANOVA (n=3 mice per group).

(B) qPCR analysis of *Il22* in the small intestines of antibiotic-treated mice administered LPS via either intragastric gavage or by intraperitoneal injection. Data for each route of administration were normalized to the corresponding vehicle control (set to 1). The y-axis is split to accommodate the substantially greater induction of *Il22* following intraperitoneal LPS administration. Significance was determined by Student's *t*-test (n=3–4 mice per group).

PAMPs, pathogen-associated molecular patterns; Abx-treated, antibiotic-treated; qPCR, quantitative real-time PCR; sm. int., small intestine; LTA, lipoteichoic acid; LPS, lipopolysaccharide; *S*. Tm, *Salmonella* Typhimurium. Each data point represents one mouse.

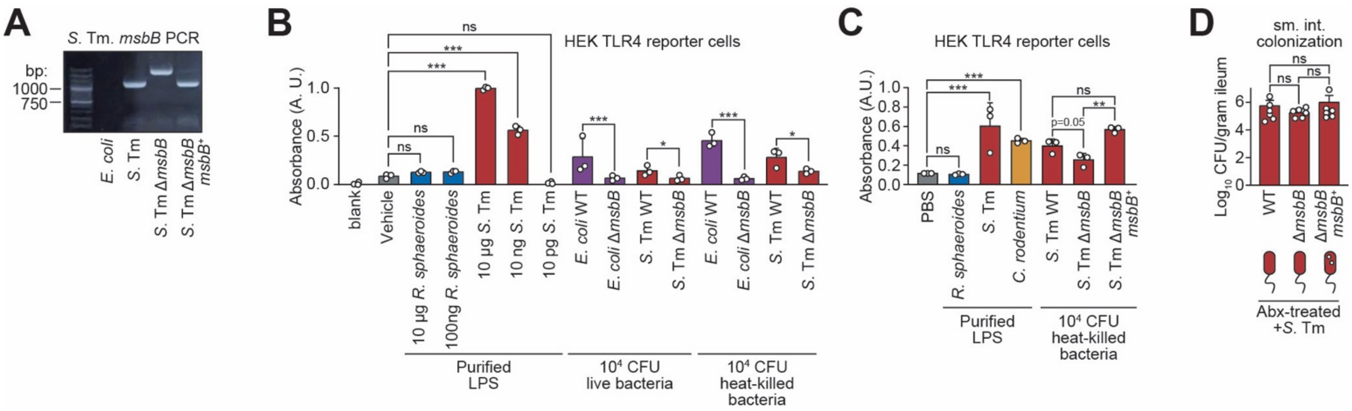


**Fig. S5. Characterization of *S*. Typhimurium and *E. coli* Δ*msbB* mutants.**

**(A)** Colony PCR gel confirming mutation of *msbB* in indicated bacterial strains. Primer sequences are listed in Table S2.

**(B)** TLR4 activation was measured in HEK reporter cells stimulated with purified LPS from *Rhodobacter sphaeroides* or *S.* Typhimurium SL1344, or with $10^4$ CFU of live or heat-killed wild-type or mutant *S.* Typhimurium IR715 or *E. coli* strains. *R. sphaeroides* expresses penta-acylated LPS that lacks the ability to stimulate TLR4 (18).

**(C)** TLR4 activation in HEK reporter cells stimulated with wild-type, Δ*msbB* mutant, or complemented *S*. Typhimurium IR715 strains.

**(D)** Bacterial burdens in distal small intestines of Abx-treated mice orally infected with wild-type, Δ*msbB* mutant, or complemented *S*. Typhimurium IR715 strains.

*S*. Tm, *Salmonella* Typhimurium; Abx-treated, antibiotic-treated; WT, wild-type; CFU, colony forming units; Uninf., uninfected; LPS, lipopolysaccharide; TLR4, Toll-like receptor 4; A.U., arbitrary units; sm. int., small intestine. Each data point represents one mouse. Significance was determined by one-way ANOVA.

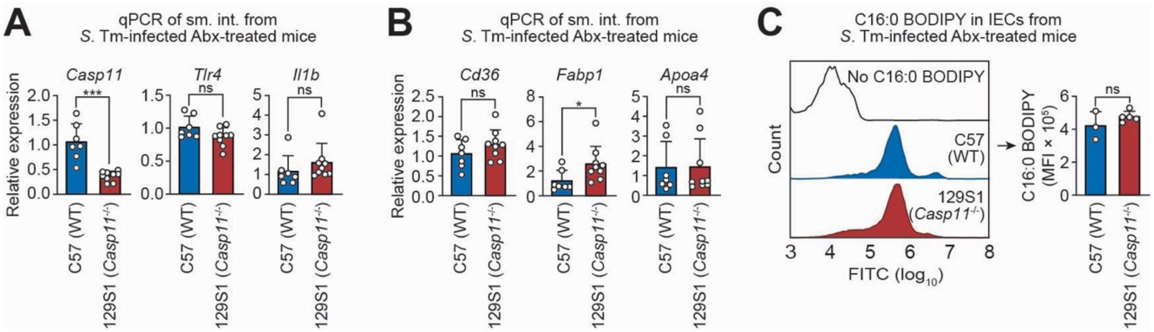


**Fig. S6. Epithelial lipid uptake is independent of Caspase-11–mediated inflammasome signaling.**

**(A)** qPCR analysis of *Casp11*, *Tlr4*, and *Il1b* expression in small intestines of wild-type C57BL/6 and Caspase-11–deficient 129S1 mice 24 h after *S*. Typhimurium infection (n=7–9 mice per group). 129S1 (129S1/SvImJ) mice lack functional Caspase-11 due to a naturally occurring loss-of-function mutation in the *Casp11* gene.

**(B)** qPCR analysis of lipid metabolism transcripts (*Cd36*, *Fabp1*, and *Apoa4*) in small intestines of antibiotic-treated C57BL/6 and 129S1 mice 24 h after *S*. Typhimurium infection (n=7–9 mice per group).

**(C)** Uptake of C16:0 BODIPY by intestinal epithelial cells (IECs) from antibiotic-treated C57BL/6 and 129S1 mice 24 h after *S*. Typhimurium SL1344 infection. Fluorescence was quantified by flow cytometry; representative histograms and quantification of median fluorescence intensity are shown (n = 3–5 mice per group).

qPCR, quantitative real-time PCR; sm. int., small intestine; *S*. Tm, *Salmonella* Typhimurium; Abx-treated, antibiotic-treated; IECs, intestinal epithelial cells; Each bar graph data point represents one mouse. Significance was determined by Student's *t*-test.

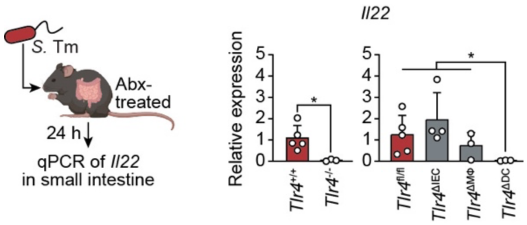


**Fig. S7. *S*. Typhimurium-induced *Il22* expression requires TLR4 signaling in dendritic cells.** Antibiotic-treated (Abx-treated) mice were orally infected with *S*. Typhimurium (*S*. Tm) for 24 hours. Small intestines from infected *Tlr4*$^{+/+}$ or *Tlr4*$^{-/-}$ mice (left) and infected *Tlr4*$^{fl/fl}$, *Tlr4*$^{\Delta IEC}$, *Tlr4*$^{\Delta M\varphi}$, or *Tlr4*$^{\Delta DC}$ mice (right) were analyzed by qPCR for *Il22* transcripts. Each bar graph data point represents one mouse (n=3–5 mice per group). *, $p < 0.05$ as determined by Student's *t*-test (left) or two-way ANOVA (right).

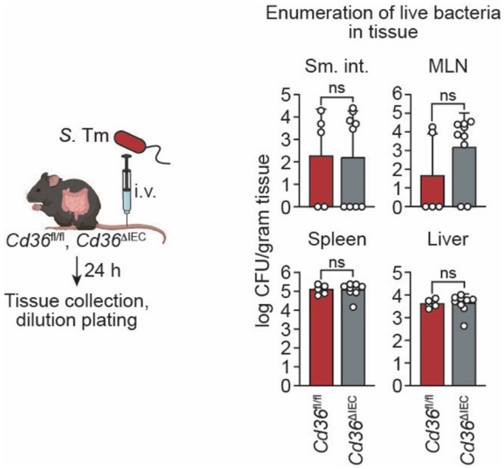


**Fig. S8. CD36 does not limit systemic *S*. Typhimurium burden following intravenous infection.**
*S*. Typhimurium SL1344 burdens in the small intestine lumen, mesenteric lymph nodes, spleen and liver 24 h after intravenous infection of *Cd36*$^{\Delta IEC}$ (n=9 mice) and *Cd36*$^{fl/fl}$ littermates (n=5 mice). Data were analyzed by Student's *t*-test. *S*. Tm, *Salmonella* Typhimurium; CFU, colony forming units; sm. int., small intestine; MLN, mesenteric lymph nodes.

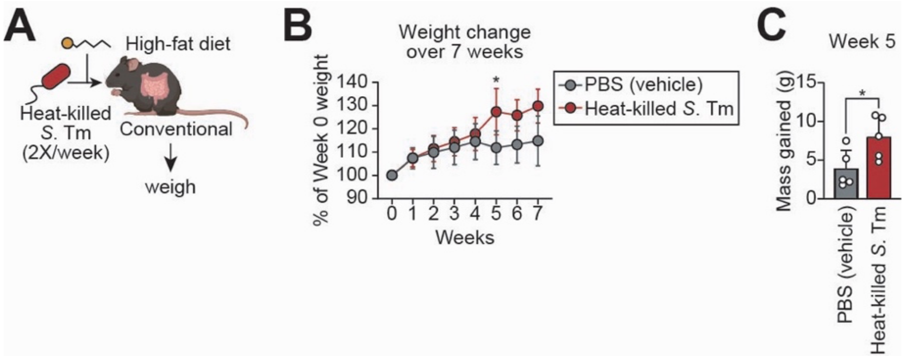


**Fig. S9. Intragastric treatment with heat-killed *S*. Typhimurium promotes weight gain in mice fed a high-fat diet.**

**(A)** Conventional C57BL/6 mice were gavaged with $10^{11}$ CFU of heat-killed wild-type *S*. Typhimurium (*S*. Tm) SL1344 or PBS (vehicle) twice a week for seven weeks. Mice were maintained on a high-fat diet for the duration of the experiment.

**(B)** Weight change over the seven-week experiment. Data were first assessed for normality using the Shapiro-Wilk test. Upon confirmation of normal distribution, percent body weight change over time was analyzed by two-way ANOVA with Sidak's post-hoc test for multiple comparisons.

**(C)** Mass gained in *S*. Typhimurium- or vehicle-treated mice at week 5. Each bar graph data point represents one mouse. Significance was determined by Student's *t*-test.

**Table S1: Lipid classes and abbreviations used in lipidomics analyses**

| Abbreviation | Lipid class |
|---|---|
| CE | Cholesterol ester |
| Hex2Cer | Dihexosylceramide |
| Cer | Ceramide |
| HexCer | Hexosylceramide |
| PC | Phosphatidylcholine |
| PI | Phosphatidylinositol |
| PS | Phosphatidylserine |
| TAG | Triacylglycerol |
| PG | Phosphatidylglycerol |
| LPC | Lysophosphatidylcholine |
| LPE | Lysophosphatidylethanolamine |
| DAG | Diacylglycerol |
| PE | Phosphatidylethanolamine |

**Table S2: Oligonucleotides used in this study**

| Description | Source | Identifier/Sequence |
|---|---|---|
| TaqMan *Apoa4* primer+probe | Thermo Scientific | Mm00431814_m1 |
| TaqMan *Fabp1* primer+probe | Thermo Scientific | Mm00444340_m1 |
| TaqMan *Cd36* primer+probe | Thermo Scientific | Mm00432403_m1 |
| TaqMan Scd1 primer+probe | Thermo Scientific | Mm00772290_m1 |
| TaqMan *Rn18s* primer+probe | Thermo Scientific | Mm03928990_g1 |
| *msbB* cloning upstream forward | IdT | 5'-ACGGCTGACGACCACACTATC |
| *msbB* cloning downstream reverse | IdT | 5'-GATTTTTCGAATTCAGGGATATACTCACTATTATTTTTT TGGTTTCCATGCTTTTC |
| *msbB* cloning upstream reverse | IdT | 5'-TGCCGATTTTCGCCATGTC |
| *msbB* cloning downstream forward | IdT | 5'-AACGTAGCGGCGCTGCCG |
| pEK3 assembly *msbB* forward | IdT | 5'-gcaaaagtGACAGAGCCGGTACGCGA |
| pEK3 assembly *msbB* reverse | IdT | 5'-tgaatgggAGCACCGTCGGTTCAACC |
| pEK3 assembly pWSK29 forward | IdT | 5'-cggtgctCCCATTCACTGCCAGAGC |
| pEK3 assembly pWSK29 reverse | IdT | 5'-ggctctgtcACTTTTGCTTTGCCACGG |
| EcN *msbB* insertion mutant forward | IdT | 5'-CTAGAGGTACCGCATGTTACAGTCAACGCGCGGC-3 |
| EcN *msbB* insertion mutant reverse | IdT | 5'-AGCTCGATATCGCATGCTTTCGCCACCCGCGCTA-3' |
